# Morphological and genomic shifts in mole-rat ‘queens’ increase fecundity but reduce skeletal integrity

**DOI:** 10.1101/2020.07.31.231266

**Authors:** Rachel A. Johnston, Philippe Vullioud, Jack Thorley, Henry Kirveslahti, Leyao Shen, Sayan Mukherjee, Courtney Karner, Tim Clutton-Brock, Jenny Tung

**Affiliations:** Department of Evolutionary Anthropology, Duke University, Durham, North Carolina 27708, USA; Department of Zoology, University of Cambridge, Cambridge CB23EJ, UK; Department of Orthopaedic Surgery, Duke Orthopaedic Cellular, Developmental, and Genome Laboratories, Duke University School of Medicine, Durham, NC 27710, USA; Department of Statistical Science, Duke University, Durham, NC 27708, USA; Department of Computer Science, Duke University, Durham, NC 27708, USA; Department of Mathematics, Duke University, Durham, NC 27708, USA; Department of Bioinformatics & Biostatistics, Duke University, Durham, NC 27708, USA; Department of Cell Biology, Duke University, Durham, NC 27710, USA; Department of Zoology and Entomology, Mammal Research Institute, University of Pretoria, 0028, Pretoria, South Africa; Department of Biology, Duke University, Durham, North Carolina 27708, USA; Duke Population Research Institute, Duke University, Durham, NC 27708, USA

## Abstract

In some mammals and many social insects, highly cooperative societies are characterized by reproductive division of labor, in which breeders and nonbreeders become behaviorally and morphologically distinct. While differences in behavior and growth between breeders and nonbreeders have been extensively described, little is known of their molecular underpinnings. Here, we investigate the consequences of breeding for skeletal morphology and gene regulation in highly cooperative Damaraland mole-rats. By experimentally assigning breeding ‘queen’ status versus nonbreeder status to age-matched littermates, we confirm that queens experience vertebral growth that likely confers advantages to fecundity. However, they also up-regulate bone resorption pathways and show reductions in femoral mass, which predicts increased vulnerability to fracture. Together, our results show that, as in eusocial insects, reproductive division of labor in mole-rats leads to gene regulatory rewiring and extensive morphological plasticity. However, in mole-rats, concentrated reproduction is also accompanied by costs to bone strength.

## Introduction

A hallmark of highly cooperative societies is reproductive division of labor. This phenomenon is best understood in eusocial insects, where environmental cues lead to reproductively and morphologically specialized castes, including one or few highly fecund “queens” [1]. These changes help support the reproductive role of queens by differentiating them from nonbreeding colony members, who forage, care for young, and engage in colony defense [1, 2]. Queens are frequently much larger than their sterile colony mates (e.g., twice as large in honey bees and Pharaoh ants [3, 4]), reflecting dramatically altered growth and development programs that are explained by changes in gene regulation [5]. Social insects thus exemplify the tight evolutionary links between reproductive division of labor, cooperative behavior, and extreme morphological plasticity.

Systems in which breeding is restricted to a single female supported by multiple nonbreeding helpers are also observed in vertebrates, including birds and mammals [6]. Here, breeding status is not determined during early development, but instead occurs in adulthood, and breeding is only achieved by those individuals who have the opportunity to transition into a reproductive role. In some species, new breeders undergo a period of accelerated growth, which may be important either for maintaining dominance or for supporting high fecundity [7–12]. While substantial gene regulatory divergence with breeding status has been described for the brain and some peripheral organs [13–15], we know little about the gene regulatory shifts responsible for breeder-associated growth patterns. Because morphological change is often crucial for ramping up offspring production, these processes are key to understanding both the basis for, and limits of, status-driven differences in growth and development.

Here, we investigate the morphological and molecular consequences of experimental transitions to breeding status in female Damaraland mole-rats (*Fukomys damarensis*). Like naked mole-rats, Damaraland mole-rats are frequently classified as ‘eusocial’ [16–18], and female helpers who transition to queens experience accelerated vertebral growth associated with increases in fecundity [9, 11]. However, it is not clear what triggers skeletal remodeling, where it is localized within the vertebral column, or whether it extends to other parts of the skeleton. Further, the gene regulatory changes that support skeletal remodeling in mole-rat queens are not known, nor are their consequences for skeletal growth potential and integrity. To address these questions, we experimentally assigned age-matched, female littermates to become queens or remain as nonbreeders and evaluated gene regulatory and morphological changes induced by the transition to queen status. Our results indicate that, as in eusocial insects, females that acquire breeding status experience substantial morphological remodeling, associated with pathway-specific changes in gene regulation. Notably, we found that queens not only experience lengthening of their lumbar vertebrae, but also show reductions in the growth potential and structural integrity of their long bones. These changes result from increased rates of bone resorption that may increase the risk of fracture, indicating that the presence of helpers does not annul the costs of reproduction to queens.

## Results

### Adaptive plasticity in the skeleton of Damaraland mole-rat queens

Adult female Damaraland mole-rats were randomly assigned to either transition to queen status (n = 12) or remain as nonbreeders (n = 18) for the duration of the experiment (Figure 1A; Supplementary Table S1). Age at assignment (mean age = 19.4 ± 4.4 s.d. months) was consistent with the age at dispersal observed in wild Damaraland mole-rats (1 – 3 years, Thorley and Clutton-Brock, unpublished data). To resolve whether skeletal changes are a function of the queen transition *per se* versus release from reproductive suppression in the natal colony, nonbreeders were either kept in their natal colonies as helpers or placed into solitary housing in the absence of a breeding queen, recapitulating extended periods of dispersal in this species [18] (n = 10 helpers and n = 8 solitaires). At the time of assignment, females assigned to the queen, helper, and solitaire treatments were statistically indistinguishable in body mass, age, or vertebral length (as measured by lumbar vertebra 5 (LV5); unpaired t-tests between all pairwise combinations of treatments: p > 0.05; Supplementary Figure S1). When possible, we assigned age-matched littermates to queen versus nonbreeding treatments (26 of 30 experimental animals were in sets of littermate sisters; Supplementary Table S1). Six non-experimental animals (1 queen and 5 nonbreeders) were also included in the sample, resulting in a total sample size of 13 breeders and 23 nonbreeders (Supplementary Table S1).

**Figure 1.**
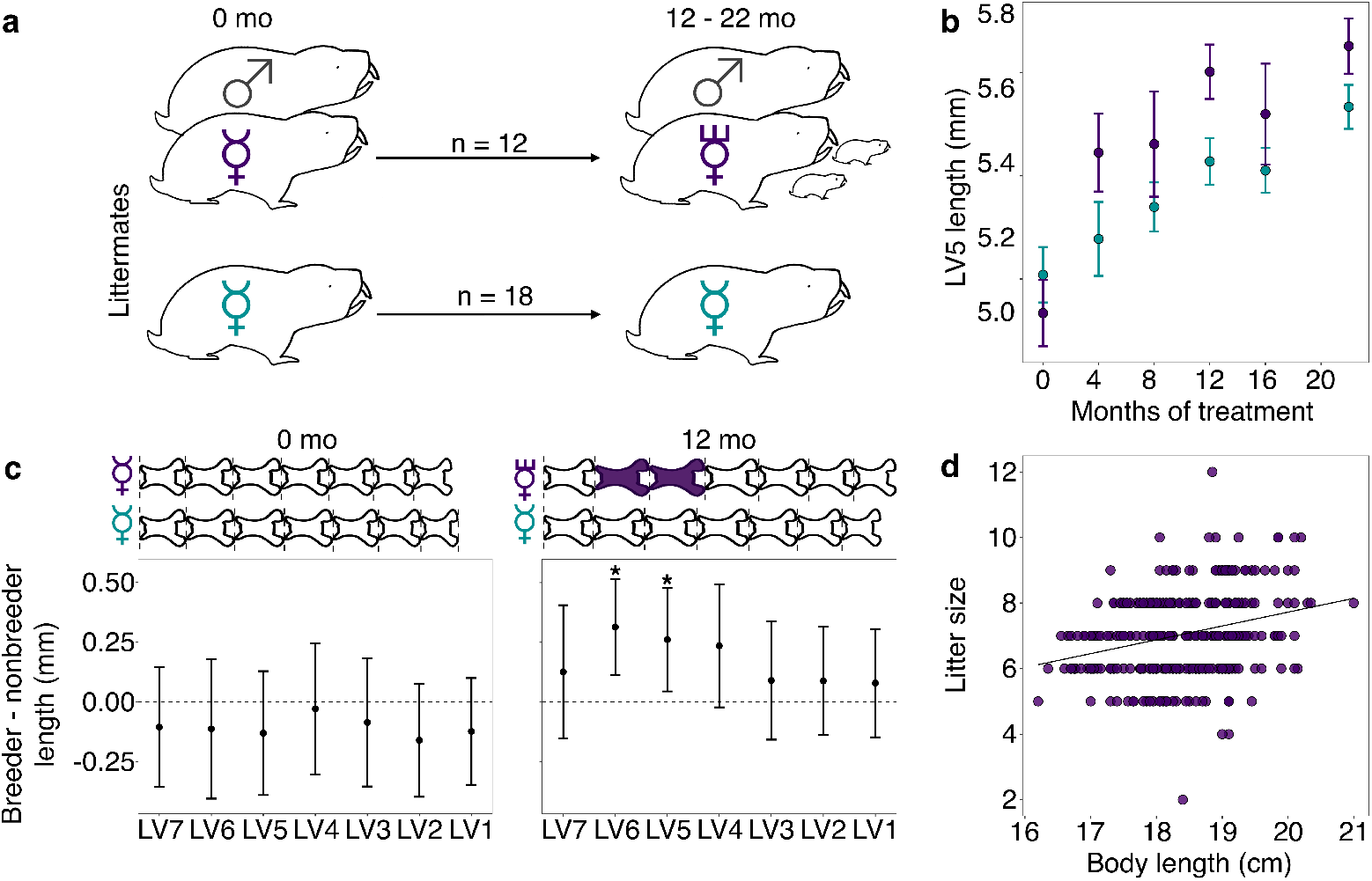
Transition to queen status leads to lumbar vertebral lengthening. **(a)** Experimental design: nonbreeding adult female (☿) littermates were randomly assigned to transition to queen status (purple 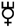) by being paired with an unrelated male (♂), or to remain in a nonbreeding treatment (cyan). Duration of treatment ranged from 12 – 22 months. **(b)** Queens show more rapid growth in lumbar vertebra 5 (LV5) in the first four months of the experiment, relative to nonbreeders (treatment by time point interaction: β = 0.078, n = 49, p = 3.47 × 10^−3^). Dots show means +/− standard errors (bars). **(c)** At the start of the experiment (0 months, left panel), the lumbar vertebrae of breeders do not differ from those of nonbreeders (unpaired t-tests, all p > 0.05). However, at 12 months (right panel), queens have longer lumbar vertebrae relative to nonbreeders (unpaired t-tests, * indicates p < 0.05). Dots show means +/− standard errors (bars). Lengths of lumbar vertebrae above the plots are scaled to indicate the mean lengths of queens (top) and nonbreeders (bottom) at each time point; vertebrae highlighted in purple are significantly longer in queens relative to nonbreeders. **(d)** Litter size is positively correlated with maternal body length in the Damaraland mole-rat colony (β = 0.353, n = 328 litters, p = 1.35 × 10^−3^).

Females assigned to the queen treatment were each transferred to a new tunnel system containing only an unrelated adult male, simulating the natural process of dispersal and new colony formation in the wild [18]. This pairing procedure, which defines the queen treatment, typically leads to immediate sexual activity and rapid activation of the reproductive axis, including initiation of ovulation and the potential for conception [19, 20]. Queens gave birth to a mean of 6.92 ± 5.57 s.d. live offspring during the 12 – 22 month follow up period, produced in a mean of 2.85 ± 1.75 s.d. litters (range: 0 – 6; Supplementary Table S1). As expected, helpers and solitaires produced no offspring, and did not differ from each other in body mass or vertebral length after the 12 – 22 month follow-up period (unpaired t-tests, all p > 0.05; Supplementary Figure S2). Because helpers and solitaires were morphologically indistinguishable, and also exhibited no differences in gene expression in our subsequent genomic assays (Supplementary Table S2), we grouped them together into a single “nonbreeder” treatment for the remainder of our analyses.

Compared to nonbreeders, queens showed rapid growth in the lumbar vertebrae in the first 12 months post-pairing (Figure 1B), especially in the vertebrae toward the caudal end of the vertebral column (LV5 and LV6). Based on longitudinal measurements, most of this differential growth was concentrated soon after the breeding status transition. Specifically, we observed a significant interaction between breeding status (queen versus nonbreeder) and post-pairing time point in the first four months of the experiment (Figure 1B; β = 0.0784, p = 3.47 × 10^−3^; n = 49 x-rays from 28 animals), but not for measurements taken in later time point intervals (4 months versus 8 months; 8 versus 12 months, all p > 0.05). Moreover, in the first four months, only queens that had already experienced pregnancy showed accelerated vertebral lengthening relative to nonbreeders (unpaired t-test; LV5 of pregnant queens vs. nonbreeders: t = −5.735, df = 16.871, p = 2.50 × 10^−5^; LV5 of queens not yet pregnant vs. nonbreeders: t = −0.789, df = 13.007, p = 0.444; n = 14 nonbreeders, 5 pregnant queens, and 2 queens not yet pregnant).

As a result of accelerated vertebral growth in queens post-transition, size differences persisted throughout the study. After 12 months, the absolute length of LV5 in queens was, on average, 4.8% longer than nonbreeders (Figure 1C; LV5: unpaired t-test, t = 2.509, df = 21.095, p = 0.020), and the absolute length of the lumbar vertebral column in queens relative to nonbreeders was 3.5% longer, although the latter difference was not significant (unpaired t-test, t = 1.945, df = 22.49, p = 0.064). Differences between queens and nonbreeders were even more apparent if lumbar vertebrae measures were scaled by zygomatic arch (head) width, as in previous studies [9, 11, 21] (LV5: 9.3% longer, unpaired t-test, t = 4.12, df = 15.135, p = 8.87 × 10^−4^; lumbar vertebral column length: 7.9% longer, unpaired t-test, t = 4.34, df = 15.37, p = 5.58 × 10^−4^). Thus, transitions to queen status induce reproductive investment, which in turn leads to organism-wide allometric changes that generate an elongated phenotype.

The elongated phenotype appears to subsequently facilitate future fecundity. Queens with longer bodies (which correlates with longer lumbar vertebrae, Pearson’s *r* = 0.856, p = 5.99 × 10^−59^; Supplementary Figure S3) had more pups per litter (Figure 1D; β = 0.353, p = 1.35 × 10^−3^, n = 328 litters from all breeding groups maintained in the same breeding facility; Supplementary Table S3). Controlling for litter size, longer queens also had larger pups: for every additional centimeter of maternal body length, pup body mass increased by 2.9% (β = 0.28, p = 0.032, n = 971 pups). Thus, the elongated queen phenotype is a strong candidate for adaptive plasticity that supports increased fertility in queen mole-rats.

### Breeding status induces gene regulatory changes in the queen mole-rat skeleton

To identify the gene regulatory changes associated with skeletal plasticity, we cultured cells enriched for bone marrow-derived mesenchymal stromal cells (bMSCs) isolated from the lumbar vertebrae (pooled LV1 – LV5) of queens and nonbreeders (n = 5 queens, 11 nonbreeders). bMSC cultures include multipotent skeletal stem cells, the precursor of the osteoblast and chondrocyte lineages responsible for bone growth. In parallel, we cultured cells enriched for bMSCs from the pooled long bones (humerus, ulna, radius, left femur, and left tibia) of the same animals, which do not show increased elongation in queens (femur at 12 months: unpaired t-test, t = −0.202, df = 19.326, p = 0.842; tibia at 12 months: unpaired t-test, t = −0.860, df = 16.759, p = 0.402). To evaluate the potential role of sex steroid hormone signaling on bone growth, we treated cells from each bone sample for 24 hours with either 10 nM estradiol or vehicle control, resulting in 47 total samples. We then performed RNA-seq on each sample to screen for genes that were systematically differentially expressed in the bone cells of queens versus nonbreeders.

Of 10,817 detectably expressed genes, 171 genes showed a significant effect of breeding status at a false discovery rate (FDR) threshold of 10% in the long bones (329 at an FDR of 20%; Supplementary Table S4). Surprisingly, no genes showed a significant effect of breeding status in the lumbar vertebrae at either FDR threshold. However, effect sizes were highly correlated between bone types overall (R^2^ = 0.75, p = 4.60 × 10^−53^), with more pronounced effects of breeding status in the long bone samples than in the lumbar vertebrae (paired t-test on breeding status effects in long bone versus vertebrae: t = 3.97, df = 317.67, p = 8.73 × 10^−5^). Importantly, breeding status-related differences were not readily attributable to differences in bone cell composition. Based on both canonical markers of bMSC lineage cells and deconvolution of the RNA-seq data using data from 27 mesenchymal or hematopoietic lineage mouse cell types, the majority cell type in both queen and nonbreeder samples was most similar to cells from the bMSC lineage [22–24] (Supplementary Figures S4 and S5). Additionally, the top three principal components summarizing estimated cell type proportions did not differ between queens and nonbreeders (all FDR > 10%, Supplementary Table S5), and we identified no cases in which the effects of breeding status on gene expression were significantly mediated by the first principal component of cell composition (p > 0.05 for all 171 queen-associated genes at 10% FDR; Supplementary Table S6).

The majority of breeding status-associated genes were up-regulated in queens (151 of 171 genes, 88%). In support of their role in skeletal plasticity, up-regulated genes were enriched for bone remodeling (log_2_[OR] = 4.07, p = 5.07 × 10^−6^), a process that involves the balanced cycle between bone formation by osteoblasts and bone resorption by osteoclasts [25] (Figure 2). Surprisingly, however, enriched pathways were specifically related to bone resorption, not formation (Supplementary Table S7), including “positive regulation of bone resorption” (Figure 2A, C; log_2_[OR] = 6.51, p = 1.55 × 10^−6^) and “superoxide anion generation,” which is involved in osteoclast activity and degradation of bone matrix (Figure 2A, C; log_2_[OR] = 5.29, p = 1.4 × 10^−5^) [26–29]. Differentially expressed genes were also enriched for immune-related processes (e.g., “cytokine secretion”, “chemotaxis”, “leukocyte activation involved in immune response”; Supplementary Table S7). These observations suggest that transitions to queen status also involve changes in immunoregulatory signaling (osteoclast cells are derived from monocytes).

**Figure 2.**
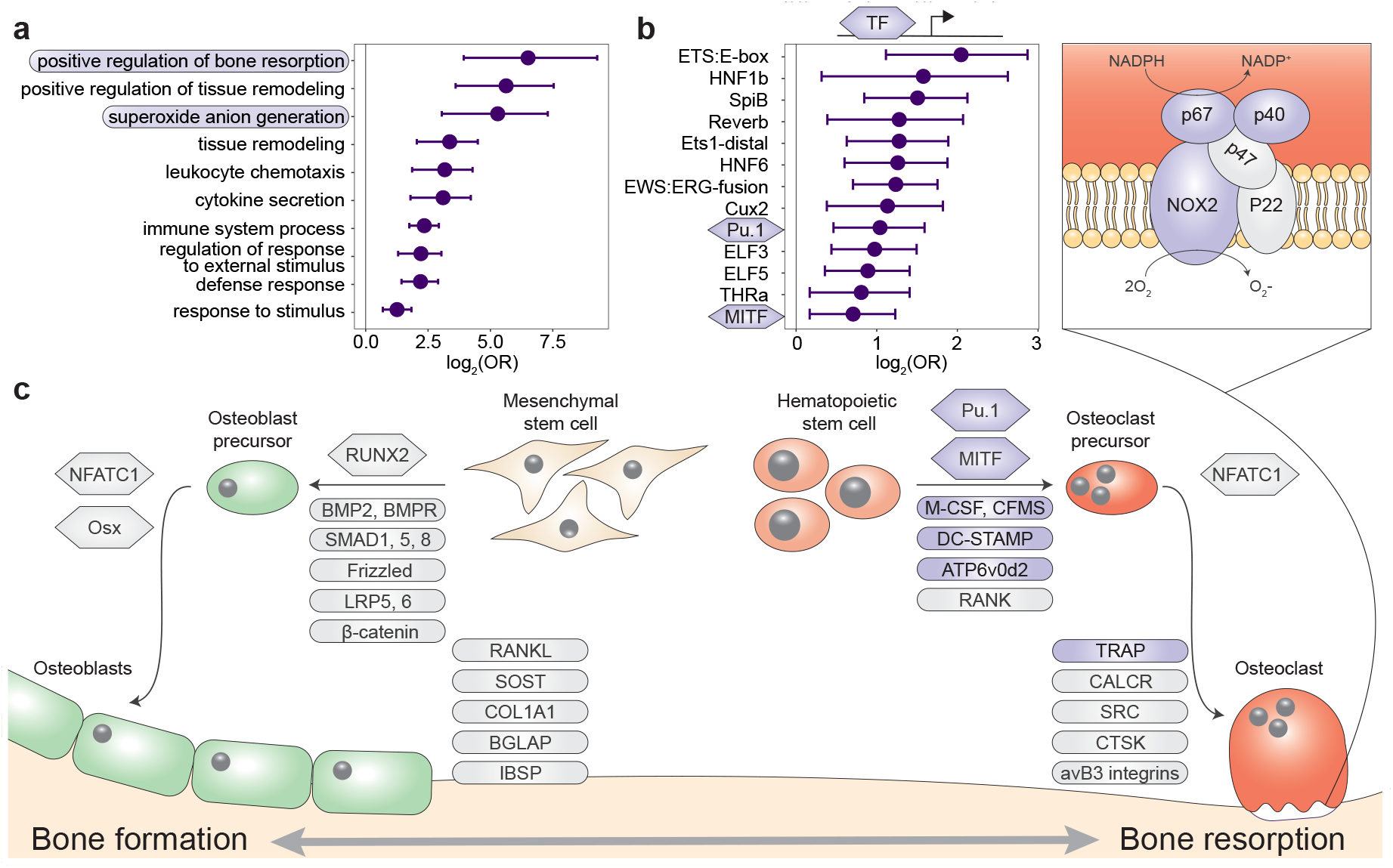
Queen status drives increased regulatory activity of bone resorption pathways. **(a)** Gene Ontology (GO) terms enriched in queen up-regulated genes, relative to the background set of all genes tested. Bars represent 95% confidence intervals. Processes highlighted in purple are also depicted in (c). Highest-level (most general) terms are shown; for full GO enrichment results, see Supplementary Table S7. **(b)** Accessible transcription factor binding site motifs enriched near queen up-regulated genes, relative to all genes tested. Bars represent 95% confidence intervals. Transcription factors highlighted in purple are also depicted in (c). **(c)** Schematic of the balance between bone formation and bone resorption, showing key regulators and markers for mesenchymal stem cell differentiation into osteoblasts and hematopoietic stem cell differentiation into osteoclasts [25, 30]. Note that not all genes or proteins in gray were tested for differential expression (e.g., because they were not annotated in the Damaraland mole-rat genome or were too lowly expressed in our sample); see Supplementary Table S4 for full set of tested genes. Queen up-regulated genes or corresponding proteins (FDR < 10%) are highlighted as purple ovals, and transcription factors with binding motifs enriched near queen up-regulated genes are highlighted as purple hexagons. Inset for osteoclasts shows the NADPH oxidase system, which generates superoxide radicals (O_2_^−^) necessary for bone resorption and is highly enriched for queen up-regulated genes (purple ovals).

Omni-ATAC-seq profiling of open chromatin regions further supports a central role for bone resorption and osteoclast activity in the queen skeleton (n = 8; Supplementary Table S8). Specifically, transcription factor binding motifs (TFBMs) located in accessible chromatin near queen up-regulated genes were enriched for PU.1 and MITF, two transcription factors that are essential for osteoclast differentiation [30] (Figure 2B, C; PU.1 log_2_[OR] = 1.041, p = 2.84 × 10^−4^; MITF log_2_[OR] = 0.707, p = 7.36 × 10^−3^; see Supplementary Table S8 for complete list of enriched TFBMs). *MITF* was also among the 151 genes that were differentially expressed between queens and nonbreeders and up-regulated in both queen long bones and lumbar vertebrae. Surprisingly, given the role of sex steroid hormones in bone growth and elevated estradiol levels in queen versus helper Damaraland mole rats [31], we observed no significant effects of estradiol treatment on gene expression in either bone type (all FDR > 10%). Queen up-regulated genes were also not in closer proximity to androgen response elements (ARE) or estrogen response elements (ERE) than expected by chance (ARE log_2_[OR] = 0.207, p = 0.627; ERE log_2_[OR] = 0.196, p = 0.652). Consistent with this observation, transcription factor footprinting analysis showed no evidence of queen-associated differences in transcription factor activity of the androgen receptor, estrogen receptor 1 (ESR1), or estrogen receptor 2 (ESR2), in either the long bones or lumbar vertebrae (all paired t-tests: p > 0.05; Supplementary Figure S6). Thus, our data point to the involvement of non-sex steroid-mediated signaling pathways in remodeling queen mole-rat bones, at least after one year post-transition.

### Extensive skeletal remodeling in queen mole-rats

The gene expression data suggest that queen status-driven changes to the skeleton extend beyond the lumbar vertebrae to the long bones. Further, they suggest that bone resorption—an important counterpoint to bone formation that is required for normal skeletal maintenance—also distinguishes breeding and nonbreeding females. To investigate this possibility, we performed high-resolution micro-computed tomography (μCT) scanning to generate 3D reconstructions of LV6, LV7, right femur, and right tibia of queens and female nonbreeders (n = 140 bones from 36 animals; Figure 3A; Supplementary Figure S7). This approach substantially increases the level of resolution for investigating breeding status-linked differences in skeletal morphology, as previous studies relied on x-ray data alone [9, 11].

**Figure 3.**
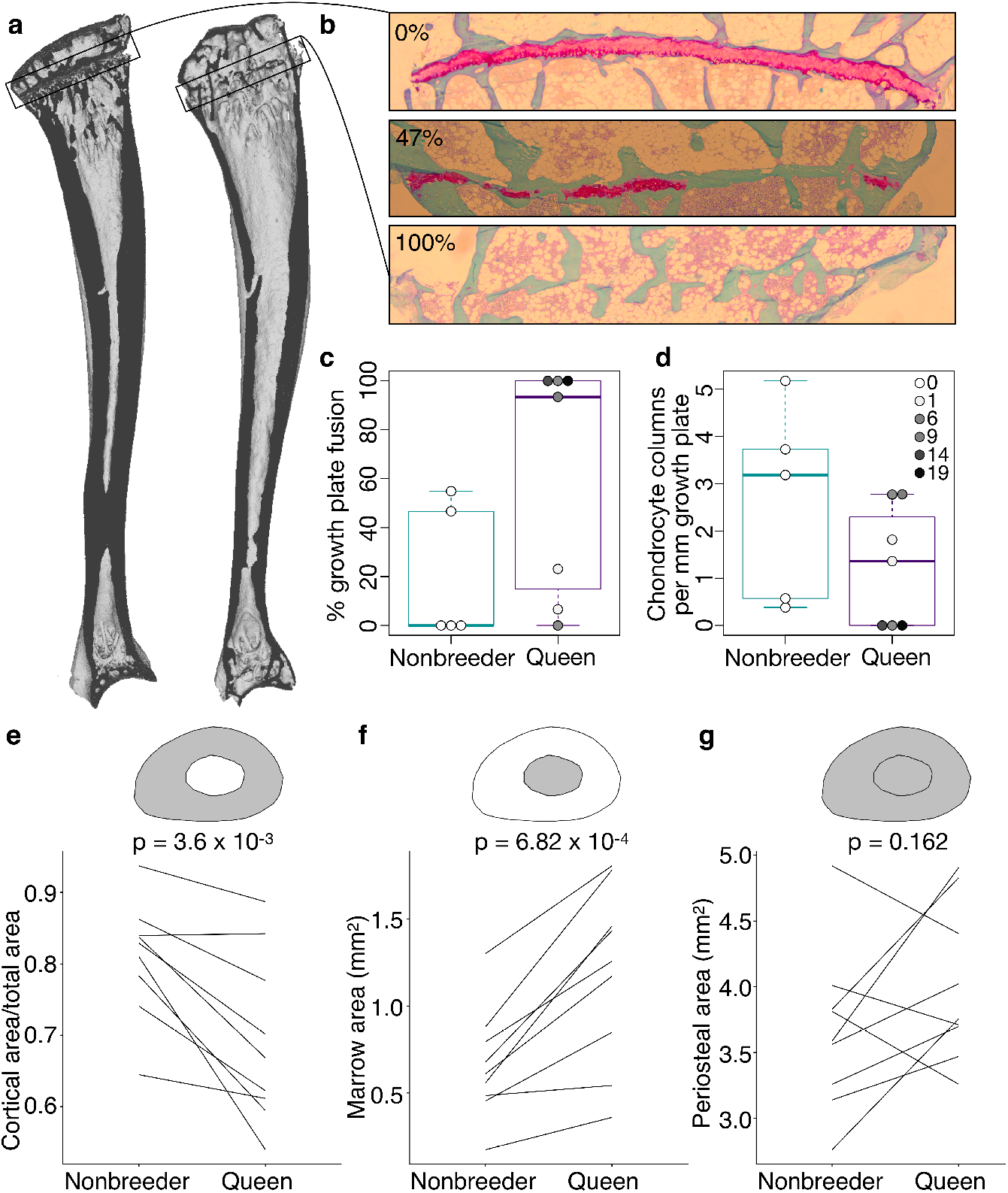
Queen status leads to reduced growth potential in the tibia and reduced cortical area at the femoral midshaft. **(a)** μCT scans of Damaraland mole-rat tibias. Boxes indicate the location of the proximal growth plate, which varies between unfused (left) to fully fused (right). **(b)**Example Safranin-O stained histological sections of the proximal tibia, in which the growth plate is unfused (top), partially fused (middle), or fully fused (bottom). Values indicate percent growth plate fusion across the width of the bone. The cartilaginous growth plate is stained deep pink, and calcified bone is stained green. **(c)** Queens, and specifically queens that gave birth to more offspring, show increased growth plate fusion (β = 0.050, p = 4.51 × 10^−3^, n = 12, controlling for age) and **(d)** decreased number of chondrocyte columns within the remaining growth plate (β = −0.132, p = 0.020, n = 12, controlling for age). Each box represents the interquartile range, with the median value depicted as a horizontal bar. Whiskers extend to the most extreme values within 1.5x of the interquartile range. In (c) and (d), dots represent individual animals, and shading indicates each animal’s total offspring number. Ages of queens and nonbreeders do not significantly differ (unpaired t-test, t = 0.489, n = 12, p = 0.644). **(e-g)** Femoral cross-sections with area highlighted in gray show the measures represented in the corresponding plots below. Each line represents an age-matched, nonbreeder and queen littermate pair. **(e)** Queens have less cortical bone (relative to the total area of the femoral midshaft cross-section), compared to their paired nonbreeding littermates (paired t-test, t = −4.07, df = 8, p = 3.60 × 10^−3^). **(f)** Queens also have enlarged marrow cavities (paired t-test, t = 5.36, df = 8, p = 6.82 × 10^−4^) but **(g)** show no difference in overall periosteal area (paired t-test, t = 1.54, df = 8, p = 0.162).

We first asked whether breeding status could be predicted from morphological differences in the 3D reconstructions. We found that it could for the lumbar vertebra, but not for the femur: by applying the smooth Euler characteristic transform [32], we were able to predict queen versus nonbreeder status in LV6 (77.8% accuracy, p = 0.01, n = 36), but not the femur (52.8% accuracy, p = 0.53, n = 36). Including only highly fecund queens (≥6 total offspring) improved predictive accuracy in the femur (70% accuracy, p = 0.12, n = 30). Although these predictions did not reach statistical significance, they raised the possibility that morphological changes in femurs become enhanced with increasing reproductive effort.

We next tested whether the transition to queen status affects the ability to continue bone lengthening. Lengthening requires the presence of a growth plate, a region of cartilage in the bone where longitudinal growth occurs through proliferation of cartilage cells (chondrocytes) (Figure 3A, B; Supplementary Figure S7). Closure of the growth plate, which indicates that bone lengthening potential has terminated, typically occurs in mammals after reaching sexual maturation, when energy begins to be invested in reproduction instead of growth [33]. To test whether the transition to queen status alters bone lengthening potential, we performed Safranin-O staining on sections of the right tibia and LV7 to visualize growth plates (Figure 3B). In the proximal tibia but not LV7, queens were less likely to have open growth plates (Figure 3C; Supplementary Figure S7 and Table S9; tibia: two-sided binomial test, p = 0.019; LV7: two-sided binomial test, p = 0.422). The increased probability of growth plate closure in the tibia of queens is linked to the number of offspring a female has produced: females with more offspring showed a higher expanse of closure across the growth plate (β = 0.050, p = 4.51 × 10^−3^, n = 12, controlling for age). This pattern may be due in part to reduced chondrocyte proliferation, as females that produced more offspring had fewer chondrocyte columns in the remaining growth plate (Figure 3D; β = −0.132, p = 0.020, n = 12, controlling for age). Thus, offspring production in queens is associated with loss of the ability to lengthen the long bones, but not the lumbar vertebrae, consistent with the importance of abdominal lengthening for supporting larger litters.

A major demand on reproductively active female mammals is a high requirement for calcium, particularly during lactation when mothers support rapid offspring bone growth. Maternal skeletons are remodeled to meet this demand, although in most mammals, these changes are not permanent (reviewed in [34]). Because of the particularly intense reproductive investment made by cooperatively breeding mole-rat queens, we therefore also evaluated the effect of queen status on trabecular and cortical bone volumes, which are thought to be important in satisfying short-term and long-term calcium demands, respectively. We found no effect of queen status on the amount of trabecular bone in the femur, tibia, LV6, or LV7 (all p > 0.05 for bone volume/total volume). However, we found that cortical bone was significantly thinner at the femoral midshaft, but not in the lumbar vertebrae, in queens compared to their nonbreeding sisters (Figure 3E; Supplementary Figure S8; femur: paired t-test of cortical area/total area, t = −4.067, df = 8, p = 3.60 × 10^−3^; LV6: paired t-test of cortical area/total area, t = −0.741, df = 6, p = 0.487). Cortical thinning in queens appears to be specifically due to increased bone resorption, which typically occurs on the endosteal (internal) surface of long bones in the marrow cavity. Indeed, queens had a larger marrow cavity (paired t-test, t = 5.355, df = 8, p = 6.82 × 10^−4^; Figure 3F) but showed no difference in periosteal area compared to their nonbreeding sisters (paired t-test, t = 1.539, df = 8, p = 0.162; Figure 3G).

Because changes in cortical bone are thought to reflect accumulated demands over long time frames, we hypothesized that cortical thinning in queens is a consequence of repeated cycles of pregnancy and lactation over time, which can occur simultaneously in Damaraland mole-rat queens. In support of this idea, we found that, within queens, the relative amount of cortical bone is not predicted by the number of pups in a queen’s most recent litter (β = −0.024, n = 13, p = 0.287), but instead by the total number of pups she produced in her lifetime. Specifically, queens who had more live births had reduced cortical bone thickness along the entire shaft of the femur (Figure 4 and Supplementary Table S10; across decile sections of the femur: all p < 0.05, controlling for mother’s litter as a random effect). Thus, cortical thinning does not commence with the transition to queen status *per se* (i.e., it is not a correlate of *achieving* breeder status), but instead appears to be a consequence of repeated investment in pregnancy and lactation. Notably, thinning is particularly marked in queens who had at least six offspring, which usually occurs by 14 months after a breeding status transition (i.e., by the third litter; Supplementary Table S10). Given that wild Damaraland mole-rat queens can maintain their status for many years [35], our results suggest that long-lived queens may experience substantial morphological change.

**Figure 4.**
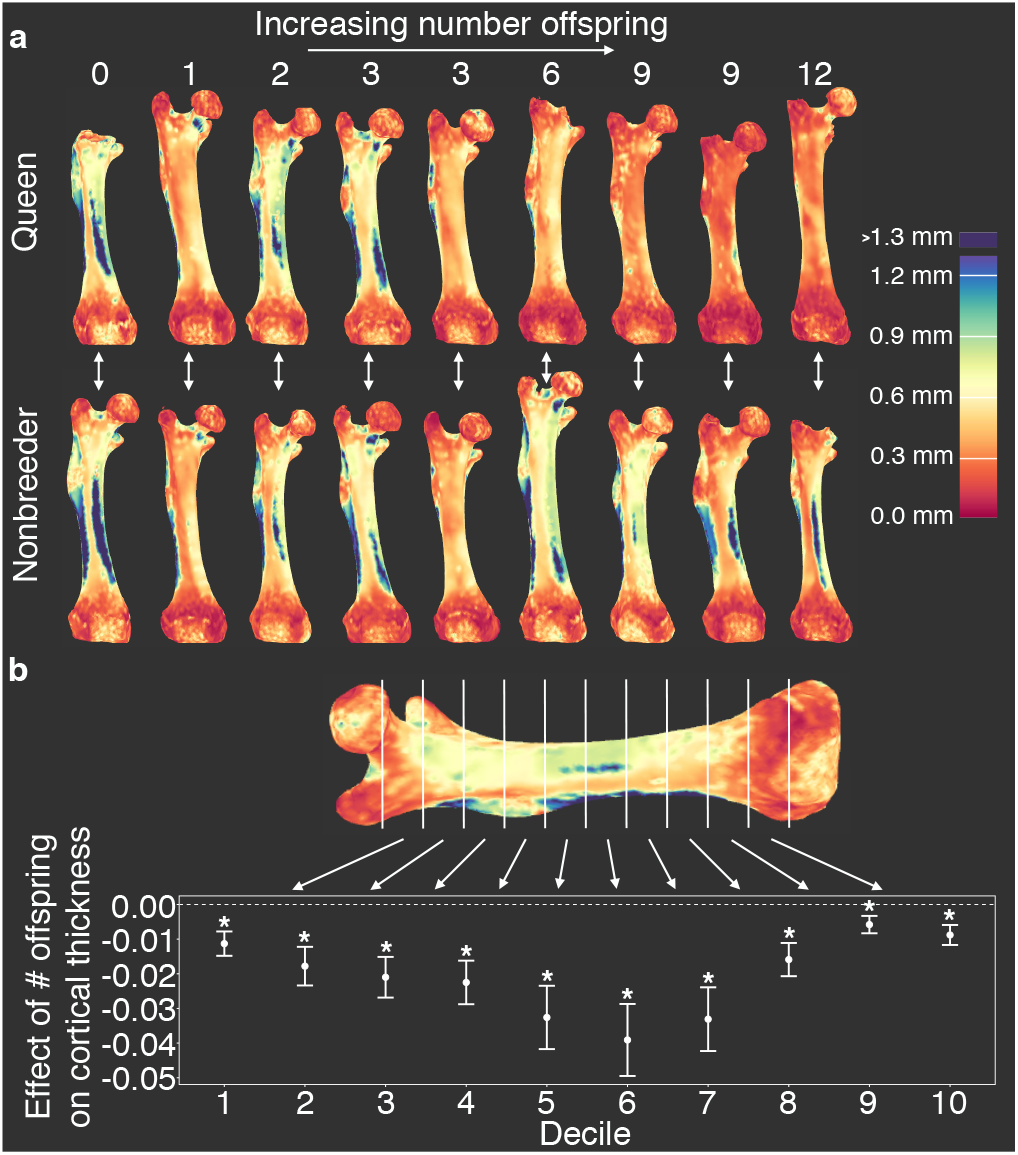
Offspring production in queens leads to cortical thinning across the femoral shaft. **(a)** Queens (top row) relative to their same-aged female nonbreeding littermates (bottom row) present thinner cortical bone across the femur, particularly in females that have many offspring (top right). Number of offspring is indicated above each femur, and vertical arrows indicate littermate pairs. **(b)** Within each decile section across the femoral shaft, number of offspring is negatively correlated with average cortical thickness (linear mixed model with littermate pair as random effect). Full results are presented in Supplementary Table S10. Asterisk indicates p < 0.05.

### Skeletal remodeling predicts increased risk of femur breakage in queens

In humans, accelerated bone resorption is a central cause of osteoporosis-related bone fragility [36]. We therefore hypothesized that cortical thinning in queen mole-rat femurs would be linked to decreased bone strength. To test this hypothesis, we calculated two key indicators of femoral structural integrity: cortical area (CA) and the minimum moment of inertia (I_min_, a predictor of resistance to bending). In nonbreeders, both are positively correlated with body mass (I_min_: R^2^ = 0.368, n = 24, p = 9.92 × 10^−4^; CA: R^2^ = 0.407, n = 24, p = 4.80 × 10^−4^). However, in queens, I_min_ is not significantly predicted by body mass (R^2^ = 0.088, n = 13, p = 0.17), but is instead a function of number of offspring produced (R^2^ = 0.283, p = 0.035). Queen CA is predicted by both offspring number and body mass, but offspring number explains almost twice the variance (offspring number R^2^ = 0.634, p = 6.83 × 10^−4^; body mass R^2^ = 0.385, p = 0.014).

To evaluate the effects of reproductive activity on the risk of bone failure, we drew on data on the relationship between CA and bone mechanical failure in a large data set of mouse femurs [37]. In this data set, CA is the best predictor of maximum load (the maximum force a bone can withstand prior to failure), and, crucially, the CA-max load relationship is highly linear (Supplementary Figure S9; R^2^ = 0.88, p = 6.64 × 10^−38^). Scaling the mole-rat CA data to mouse suggests that transitions to queen significantly increase the risk of bone failure (Figure 5; hazard ratio (95% confidence interval) = 2.67 (1.20, 5.93), n = 36, p = 0.016). Similar to growth potential and cortical thinning, this effect is driven by highly fertile queens, such that those who had at least six offspring showed the highest predicted risk of bone failure (Figure 5; queens with ≥ 6 offspring relative to nonbreeders: HR = 3.81 (1.47, 9.83), n = 30, p = 0.006). The risk of bone failure is thus predicted to increase by 21% for each additional pup (HR = 1.21 (1.10, 1.33), n = 36, p = 8.85 × 10^−5^).

**Figure 5.**
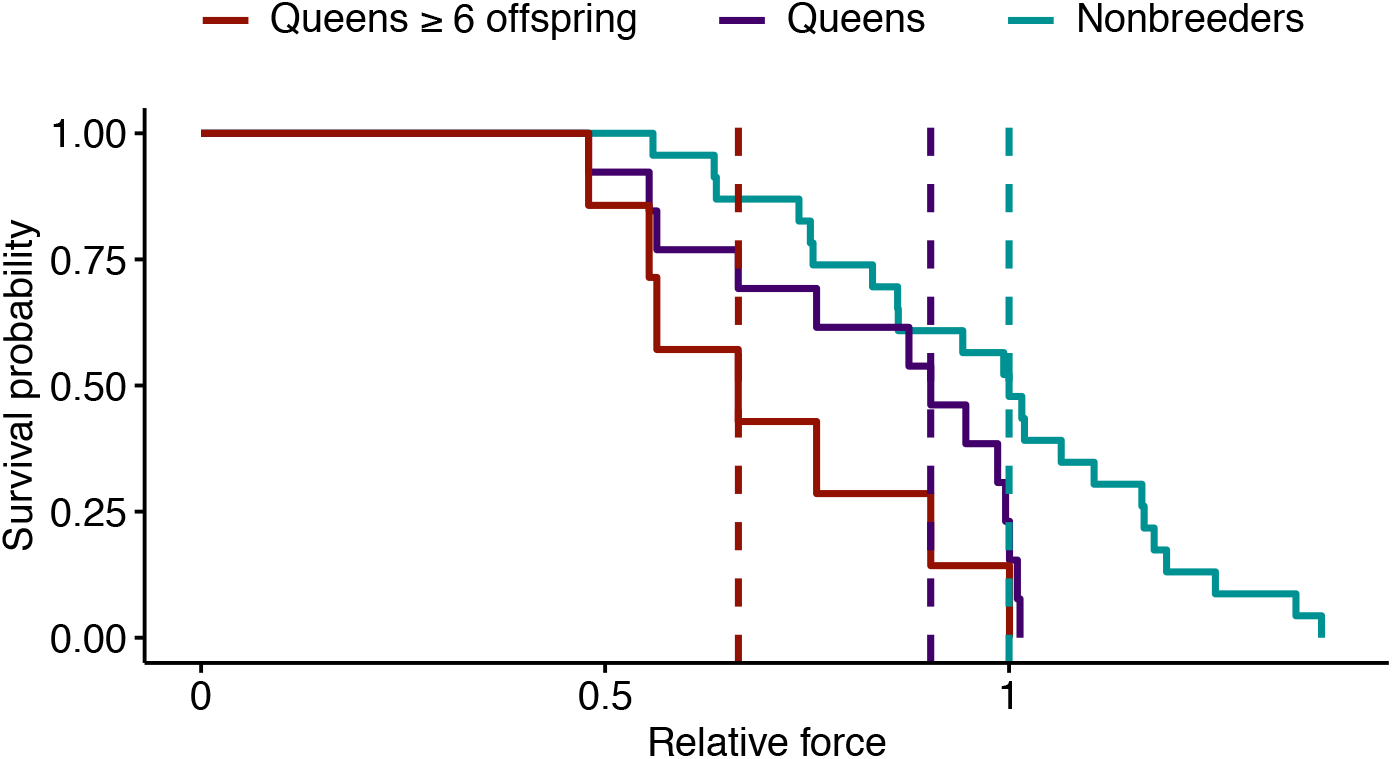
Effect of reproductive status on the probability of bone failure. Survival curves for femurs from nonbreeders versus queens (Wald test, p = 0.02, n = 36) and versus queens with ≥ 6 offspring (Wald test, p = 0.006, n = 30), based on predictions from the midshaft cortical area and data from [37]. Vertical dashed lines indicate group medians, with the median failure time for nonbreeders fixed at a value of 1.

## Discussion

Our results demonstrate that transitions to breeding status in Damaraland mole-rat queens lead to a cascade of skeletal changes linked to shifts in gene regulation. The vertebral lengthening observed in Damaraland mole-rat queens is concordant with previous reports of vertebral lengthening in both Damaraland mole-rats [11] and naked mole-rats [9]. Like naked mole-rats, our analyses show that most growth is concentrated soon after the breeding status transition, especially in connection with the first post-transition pregnancies [21, 38]. However, our findings also suggest subtle differences: for instance, while the growth phenotype in naked mole-rats occurs at the cranial end of the lumbar vertebrae [38], it is concentrated at the caudal end of the vertebral column in Damaraland mole-rats. Given that Damaraland mole-rats and naked mole-rats independently evolved a similar, highly cooperative social structure [18, 39], this difference suggests potential convergent evolution of the vertebral lengthening phenotype in queens, presumably in response to the selection pressure for increased fertility.

In addition to previously described vertebral growth, we found that queen Damaraland mole-rats lose bone lengthening potential in the long bones and develop thinner femurs that are predicted to be more prone to mechanical failure. Moreover, gene expression levels in queens reflect a signature of bone resorption, rather than bone growth, at the time of sampling, which occurred 1 – 2 years post-transition. The molecular signature of bone resorption temporally aligns with changes in morphology, in which accelerated vertebral growth primarily occurs during a female’s first pregnancy, whereas cortical thinning in the long bones are a function of repeated cycles of offspring production. Thus, queens quickly progress from traits typically associated with pre-reproductive and pubertal growth in mammals (e.g., body elongation), to traits typically linked to aging (e.g., marrow cavity expansion and cortical thinning).

The complex pattern of bone growth and bone resorption in queens likely involves multiple regulatory mechanisms. Because estrogen is known to impact bone growth and maintenance [40, 41], and estrogen levels are higher in mole-rat queens relative to nonbreeding females [31], we hypothesized that queens and nonbreeders would differ in their response to estradiol in bone marrow-derived cells. Surprisingly, we observed no gene expression response to estradiol treatment. By itself, this result could be a function of the specific concentration or duration of estradiol treatment we applied. However, we also observed no enrichment for estrogen receptor binding motifs near queen up-regulated genes, and no evidence that estrogen or androgen receptor binding sites are differentially bound in cells from queens versus non-breeders. Thus, our results suggest a role for other, as-yet unknown signaling pathways in the queen-specific signature of long bone cortical resorption (although it does not exclude the importance of estrogen signaling in other phenotypes, such as bone elongation or growth plate closure, [42]).

Bone loss in Damaraland mole-rat queens may be an extreme of the typical mammalian pattern of bone remodeling, in which bone mineral density decreases during pregnancy and lactation, but recovers once offspring are weaned [34]. Thinning in mole-rats may be sustained, however, because queens begin gestating soon after lactating for the previous litter, leaving little to no time for recovery. One possible reason that this fast rate of breeding is achievable is that queens in colonies with more helpers work less and rest more [43], consistent with studies in other cooperative mammals that show that helpers alleviate breeding-associated loss of condition in queens [44]. Paradoxically, helpers might not only help offset costs of, but also contribute to, decreased bone mass in queens, given that large numbers of helpers are themselves produced via high queen fecundity, and reduced physical activity can also lead to decreases in bone mass [45].

The extent to which helpers reduce the costs of breeding to queens may also differ between species depending on the relative numbers of helpers to breeders. For example, in eusocial insects, large colonies and the high ratio of helpers to queens reduce the costs of reproduction to queens to very low levels [1, 2]. Similarly, in naked mole-rats (where colonies can include hundreds of animals, compared to dozens in Damaraland mole-rat colonies [17, 18, 46]), a small sample of queens (n = 6) suggests increased rather than decreased femoral cortical thickness relative to age-matched nonbreeders [47]. Testing how the costs and benefits of reproduction are resolved across different levels of cooperativity, including the molecular mechanisms that mediate these differences, is an important next step towards understanding the evolution of cooperative breeding in mammals.

Finally, despite frequent analogies between Damaraland mole-rats and eusocial insects [18, 46, 48], our results suggest some key points of differences. Specifically, while abdominal lengthening allows queen mole-rats to increase fecundity per reproductive effort, loss of cortical bone in the femur is unlikely to directly benefit either fertility or survival. Instead, it reflects the cumulative burden of continuous cycles of pregnancy and lactation [34]. Thus, unlike eusocial insect queens [49, 50], Damaraland mole-rat queens incur morphological costs to concentrated reproduction in addition to morphological changes that facilitate increased fitness. How these costs translate into fertility or survival outcomes in natural populations remains a fascinating, unanswered question.

## Materials and Methods

### Study system and experimental design

Damaraland mole-rats (*Fukomys damarensis*) were maintained in a captive colony at the Kuruman River Reserve in the Northern Cape Province of South Africa, within the species’ natural range. Only animals born in captivity, with known birthdates and litter composition, were used in this study, so that exact ages were known. Animals were maintained in artificial tunnel systems built from PVC pipes with compartments for a nest-box and waste-box and transparent windows to allow behavioral observation [51]. Animals were fed *ad libitum* with sweet potatoes and cucumbers.

Adult females (> 1 year) from 16 natal colonies were randomly assigned to be either nonbreeders or queens, such that females assigned to queen status had age-matched littermates, where possible, who were assigned to the nonbreeding condition. To distinguish the effects of queen status from release from reproductive suppression, nonbreeders were either maintained in their natal colony as helpers or maintained alone, which models the social condition experienced by dispersing females. Females assigned to the breeder condition were transferred to a new tunnel system with an unrelated male from a separate social group. Nine new breeding females, 6 helpers, and 8 solitaire females (age-matched littermates where possible; Supplementary Table S1) were set up in December 2015 – July 2016 [11]. With one exception (animal G10F026), animals maintained their breeding status for 14 – 22 months before sample collection. One queen and five helpers that were siblings, but not age-matched littermates, of experimental animals were also included in sample collection. To increase the final sample size, an additional 4 breeding colonies, matched against 4 age-matched littermate helpers, were formed in October 2017 and followed for 11-12 months (Supplementary Table S1). One queen died before sample collection, and one non-experimental helper was euthanized during the course of the study and included in sample collection. The final sample size included 13 queens, 15 helpers, and 8 solitaire females.

### X-ray data

For a subset of study subjects, full body X-rays were taken using the Gierth TR 90/20 battery-operated generator unit with portable Leonardo DR Mini plate (OR Technology, Rostock, Germany) every two months during the first 12 months of the experiment and at the time of sacrifice. From these X-rays, an experimenter blind to animal breeding status measured the length of each lumbar vertebra (from vertebra 1 to 7), the right femur, the right tibia, body length, and the width of the zygomatic arch using ImageJ [52]. The caudal-most lumbar vertebra was labeled as LV7. We tested for an effect of breeding status on LV5 using a linear mixed model in which post-pairing time point, breeding status, and the interaction of time point by breeding status were modeled as fixed effects and animal ID as a random effect.

### Effects of queen body length on fertility

To test the effect of maternal body length on litter size and pup size, we used body length measurements obtained during routine colony monitoring of all queens maintained in the captive colony (i.e., not restricted to experimental animals). Following [11], we used body length measurements obtained nearest to, and no more than 90 days from, the date of parturition. The resulting dataset included 328 litters (971 pups) from 76 mothers, which represents a 76% increase over an earlier analysis of this relationship in [11]. We fit two linear mixed effects models. In the first model, we modeled litter size as a function of maternal body length, controlling for whether the litter was the female’s first litter, and included maternal ID as a random effect. In the second model, we modeled pup mass as a function of maternal body length, controlling for litter size and whether the litter was the female’s first litter as fixed effects, and maternal ID and litter ID as random effects.

### Sample collection and cell culture from lumbar vertebrae and long bones

Animals were deeply anesthetized with isoflurane and sacrificed with decapitation following USGS National Wildlife Health Center guidelines and under approval from the Animal Ethics Committee of the University of Pretoria. Immediately upon sacrifice, the lumbar vertebrae and long bones were dissected, and attached muscle tissue was removed with forceps. Lumbar vertebrae 6 and 7 and the right femur and tibia were collected into 50% ethanol for 24 hours, then transferred to 70% ethanol and stored at 4° C for μCT scans and histochemistry.

To isolate bone cells for culture, lumbar vertebrae 1 – 5 were incubated in 2% Collagenase P (Roche, Switzerland) for 30 minutes at 30° C. Each bone was then cut in half and transferred to a 1.5 ml microcentrifuge tube containing a G-Tube microcentrifuge tube (VWR, Radnor, PA, USA) that had been punctured at the bottom with a 15 gauge needle. Tubes were spun at 3,000 RCF for 5 seconds, allowing the marrow to collect into the 1.5 mL microcentrifuge tube. Cell pellets were resuspended in red blood cell lysis buffer, pooled, and incubated for 3 minutes at room temperature. 10 ml bMSC media (MEM-alpha [ThermoFisher, Waltham, MA, USA] + 15% fetal bovine serum [Hyclone, Logan, UT, USA] + 1% penicillin/streptomycin + 2 ng/ml recombinant human fibroblast growth factor-basic [Biocam, Centurion, Gauteng, South Africa] + 10 nM ROCK inhibitor Y-27632 [RI; Cayman Chemical, Ann Arbor, MI, USA]) was added to stop lysis, and the tubes were spun for 5 minutes at 300 RCF. The cell pellet was resuspended in 1 ml bMSC media and strained through a 70 μm cell strainer. Cells were plated at 1.6 × 10^5^ cells per cm^2^. The long bones (excluding right femur and tibia) were processed to enrich for bMSCs following the same procedure, but without incubation in Collagenase P. Cells were cultured at 37**°** C and 5% CO_2_. Twenty-four hours post plating, plates were carefully washed three times with 1x PBS and supplied with fresh media to remove non-adherent cells. Once bMSC clusters were visible (2 – 9 days post plating), plates were fed bMSC media without RI or fed bMSC media without RI + 10 nM estradiol (E2). Twenty-four hours later, cells were collected into buffer RLT and stored at −80**°** C. Samples were shipped on dry ice to Duke University for RNA extraction using the Qiagen RNeasy Micro Kit. RNA-Seq libraries were generated using the NEBNext Single Cell/Low Input RNA Library Prep Kit for Illumina.

### Gene expression analysis

RNA-Seq libraries were sequenced on an Illumina HiSeq 4000 (100 base pair single end reads) to a mean coverage of 16.1 ± 3.9 s.d. million reads. Reads were trimmed with *cutadapt* version 2.3 [53] (parameters: −q 20 −e 0.2 --times 5 --overlap 2 -a AGATCGGAAGAGC -a “T” --minimum-length=20). Trimmed reads were then mapped to the Damaraland mole-rat v1.0 genome [54] (DMR_v1.0) using two pass mapping with STAR [55]. Only uniquely mapped reads were retained. HTseq [56] was used to quantify read counts mapping to genes (using the v1.0.92 gtf file from Ensembl; we extended the genomic coordinates of the *SERPINE1* gene by 2000 basepairs in both directions due to very high expression directly adjacent to the annotated coordinates). We transformed read counts to transcripts per million (TPM) [57], and retained only genes with TPM ≥ 2 in at least 25% of samples. We performed voom normalization [58] on the raw counts, using normalization factors produced by the trimmed mean of M-values (TMM) method [59] in DESeq [60]. We used the *limma* [61] function *lmFit* to regress out the proportion of uniquely mapped reads in genes (which controls for efficiency of mRNA selection during RNA-Seq library preparation) and animal natal colony (which controls for littermate sets and date of sacrifice) to obtain normalized, batch-corrected gene expression values for downstream analysis. We used the mixed effects model approach in *emmreml* [62] to estimate, for each gene, the effect of breeding status on gene expression within lumbar vertebrae and within long bones using the following model:

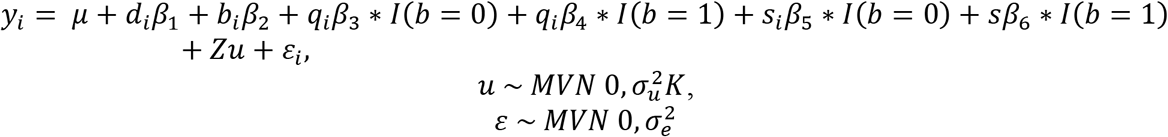

where *y* is the vector of gene expression levels for *n* = 47 samples (indexed by *i*); μ is the intercept; *d* is the number of days in culture and *β*_1_ its effect size; *b* is bone type (i.e., long bones or lumbar vertebrae) and *β*_2_ its effect size; and *q* is a 0/1 variable representing breeder status and *β*_3_ and *β*_4_ its effect size in long bones and lumbar vertebrae, respectively. *I* is an indicator variable for bone type (0 = long bone; 1 = lumbar vertebrae). *s* is a 0/1 variable representing whether the cells were cultured with estradiol and *β*_5_ and *β*_6_ are its effect sizes in long bones and lumbar vertebrae, respectively. Z is an incidence matrix that maps samples to animal ID to take into account repeated sampling from the same individual, and *u* is a random effect term that controls for relatedness. *K* is an *m* by *m* matrix of pairwise relatedness estimates (derived from the genotype data, described below) between all *m* animals. *ε* is the residual error, 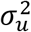 is the genetic variance component, and 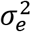 is the environmental variance component. We also ran an identical model but with an additional fixed effect of solitaire status in long bones and in lumbar vertebrae, to test for a difference in gene expression between helpers and solitaires. To control for multiple testing, we calculated the false discovery rate following Storey and Tibshirani [63] using an empirical null distribution derived from 100 permutations of each variable of interest.

We used g:profiler [64] to perform gene ontology enrichment analysis of the genes up-regulated with queen status in lumbar vertebrae and long bones (151 of 171 genes significantly associated with queen status at a 10% FDR). All genes in the original analysis set were used as the background gene set. We set both the minimum size of the functional category and the minimum size of the query/term intersection to 3. Finally, we retained categories that passed a Bonferroni corrected p-value of 0.05.

### Genotyping to estimate relatedness

To control for relatedness when modeling the gene expression data, we performed single nucleotide polymorphism (SNP) genotyping of the RNA-Seq data using the Genome Analysis Toolkit [65] (GATK). We used the SplitNCigarReads function on the trimmed, uniquely mapped reads and performed GATK indel realignment. Base recalibration was performed by using all SNPs with GQ ≥ 4 in an initial UnifiedGenotyper run on the full data set as a reference. Genotypes were called on the recalibrated bam files using HaplotypeCaller. Variants were filtered with the following GATK VariantFiltration parameters: QUAL < 100.0, QD < 2.0, MQ < 35, FS > 30, HaplotypeScore > 13, MQRankSum < −12.5, ReadPosRankSum < −8. Variants were further filtered with *vcftools* [66] to only retain biallelic SNPs in Hardy-Weinberg equilibrium (p > 0.05) with minor allele frequency ≥ 0.1, minimum mean depth of 5, max missing count of 2, and minimum GQ of 99. Finally, SNPs were thinned to a distance of 10 kb basepairs, resulting in a final dataset of 1,965 stringently filtered biallelic SNPs. Missing values were imputed using *beagle* [67], and the resulting vcf file was used to create a kinship matrix using *vcftools* [66]. Values of the kinship matrix were confirmed to be higher in known siblings compared to non-siblings (unpaired t-test, t = 27.939, p = 2.23 × 10^−12^; means = 0.513 and −0.097). Two pairs of siblings were found to have different fathers (G1F022 and G1F025; G4F020 and G4F019).

### Cell type heterogeneity

Although selection for adherent cells from bone marrow enriches for bMSCs, other cell types are also present [68]. To assess whether cell type heterogeneity accounts for queen-associated differential expression, we used CIBERSORT to deconvolve the proportion of component cell types from the RNA-Seq data [24]. We trained CIBERSORT on a data set of quantile normalized gene expression values from mouse purified primary cell populations [23]. Specifically, we subset the training data to 27 purified cell populations of mesenchymal or hematopoietic origin (Figure S3) and to genes that were included in our mole-rat gene expression dataset. We then predicted the composition of the cells that contributed to the mole-rat quantile normalized gene expression data set, for each sample.

To test whether cell type heterogeneity was significantly explained by queen status, we also modeled cell type proportion (as summarized by the first principal component of CIBERSORT-estimated proportions for all 27 potential cell types; PC1 explains 50.9% of the overall variation) following the same method used for gene-by-gene expression analysis but with PC1 included as an explanatory variable. We then performed mediation analysis on each of the 171 genes that showed a significant effect of breeding status at FDR < 10%. To do so, we first estimated the indirect effect of breeding status on gene expression through the mediating variable (CIBERSORT PC1). The indirect effect of breeding status through CIBERSORT PC1 was estimated by calculating the difference in the effect of breeding status between two models: one model that did not include the mediator (i.e., *β*_3_ and *β*_4_ from equation 1 above) and the same model with the addition of the mediating variable. We performed 1000 iterations of bootstrap resampling to obtain the 95% confidence interval for the indirect effect, and considered an indirect effect for a gene significant if the 95% interval did not overlap 0.

### ATAC-seq data and transcription factor binding site analysis

To investigate whether differentially expressed genes were associated with accessible binding motifs for specific transcription factors, we generated Omni-ATAC-seq data to profile regions of open chromatin [69, 70]. We performed Omni-ATAC-seq on both lumbar vertebrae bMSCs and long bone bMSCs from two female nonbreeding and two queen mole-rats (n = 8 libraries total), following the published protocol [70] with the following modifications: 5,000 cells were centrifuged at 500 RCF for 5 minutes at 4°C. The pellet was resuspended in 50 ul transposition mix (25 ul 2xTD buffer, 16.5 ul PBS, 6.75 ul water, 1 ul 10% NP40, 1 ul 10% Tween-20, 1 ul 1% digitonin, and 0.25 ul Tn5 transposase). The reaction was incubated at 37°C for 30 minutes without mixing, followed by a 1.5x Ampure bead cleanup. Omni-ATAC libraries were sequenced on a NovaSeq 6000 as 100 basepair paired-end reads to a mean coverage (± SD) of 26.9 (±4.4) million reads (range: 16.8 – 38.3). Reads were trimmed with Trim Galore! [71] to remove adapter sequence and low quality basepairs (Phred score < 20; reads ≥ 25 bp). Read pairs were mapped to the DMR_v1.0 genome using *bwa-mem* [72] with default settings. Only uniquely mapped reads were retained. The alignment bam files for each treatment (breeding or nonbreeding) were merged, and open chromatin regions were identified using MACS2 v2.1.2 [73] with the following parameters: “-nomodel-keep-dup all −q 0.05”. We combined open chromatin peaks with regions in the DMRv1.0 genome that match sequences of vertebrate transcription factor binding site motifs, using motifs defined in the HOMER database [74]. We used Fisher’s exact tests (using a p-value threshold of 0.01) to test if transcription factor binding motifs belonging to the same transcription factor were enriched in open chromatin regions within 2,000 bp of the 5’ most transcription start site of queen up-regulated genes.

To compare genome-wide signatures of DNA-transcription factor binding for androgen receptor (AR), estrogen receptor 1 (ESR1), and estrogen receptor 2 (ESR2), we characterized transcription factor footprints in queens and nonbreeders, in both the lumbar vertebra and long bones, using HINT-ATAC from the Regulatory Genomics Toolbox (RGT) with default parameters [75]. We focused on the subset of peak regions called using MACS2 [73]. We identified TF footprints by merging reads within each bone type-breeding status combination and calling footprints on the combined data. For each bone type, we then created a meta-footprint set by merging the respective footprint calls across queens and nonbreeders using the *bedtools* function *merge* [76]. We identified transcription factor motifs in the *DMR_v1.0* genome that fell within meta-footprints, based on the JASPAR CORE Vertebrates set of curated position frequency matrices [77]. Finally, we tested for differential footprints of AR, ESR1, and ESR2 binding using the RGT *differential* function, using the activity score metric described in [75] and default parameters.

### Micro-CT scans and analysis

We performed μCT scans of LV6, LV7, right femur, and right tibia using a VivaCT 80 scanner (Scanco Medical AG, Brüttisellen, Switzerland) set at 55 kVp and 145 μA, with voxel size 10.4 μm. Trabecular bone was quantified from the 100 μCT slices below the proximal tibia growth plate, the 100 μCT slices above the distal femur growth plate, and the 100 μCT slices medial to the caudal growth plate of LV6. To obtain midshaft cross-sections of the femur, tibia, and LV6, we first reduced each bone mesh to 100,000 faces using Avizo Lite version 9.7.0. Mesh files from the same bone type were auto-aligned using Auto3dgm [78] in Matlab. Aligned mesh files were then back scaled to their original sizes in Matlab, and the midshaft cross-section was generated using Rhinoceros version 6. The MomentMacro plugin in ImageJ was used to calculate bone area and moment of inertia.

### Classification of breeding status from bone shape

To predict breeding status from bone shape, we first applied the smooth Euler characteristic transform [32] to the aligned right femur and LV6. We then performed leave-one-out predictions, running each bone type separately, using the linear kernel and c-classification with the support vector machine (SVM) implemented by the R package *e1071* [79]. The SVM classifier was equipped with 1:100,000 weighting to achieve class balanced predictions. The empirical p-values were estimated for each bone type by running 100 permutations of the queen/nonbreeder labels [80].

### Histochemistry

For a subset of individuals (Supplementary Table S9), the tibia and LV7 were plasticized, sectioned, and stained with Safranin O by the Washington University Musculoskeletal Research Center. The proportion of the tibia proximal growth plate that was fused, and the mean proportion of the LV7 cranial and caudal growth plates that were fused, were measured in ImageJ from Safranin O stained sections. To quantify growth plate activity, we calculated the number of chondrocyte columns (defined as linear stacks of at least three chondrocytes) controlling for length of open growth plate. For each bone type (tibia and LV7), we ran two models: proportion growth plate fusion or chondrocyte columns per mm growth plate as the dependent variable, and number offspring born and age as the independent variables.

### Cortical thickness across the femur

We used Stradview [81, 82] on dicom images from the μCT scans to measure and visualize, in an automated manner, cortical thickness across the surface of the femur. Bone surface was defined in Stradview by thresholding pixel intensity and contouring the bone at every 14 sections, with the following parameters: resolution = medium, smoothing = standard, strength = very low, contour accuracy = 6. To measure cortical thickness, we used the auto threshold method in Stradview, with line width set to 5, smooth set to 1, and line length set to 3 mm. The smoothed thickness values of each femur were then registered (i.e., mapped) to a single “canonical” femur surface (mole-rat GRF002) using wxRegSurf v18 (http://mi.eng.cam.ac.uk/~ahg/wxRegSurf/). We sectioned the cortical thickness values into deciles according to location along the length of the femur. The top and bottom deciles were removed, because cortical and trabecular bone towards the ends of the femur could not be easily differentiated by the automated method. Deciles were then recreated for the remaining length of the bone (i.e., the central 80%). From each bone decile, we estimated cortical thickness as the mean of all cortical thickness measures within that interval. For each decile across animals, we used a linear mixed model to model cortical thickness as a function of breeding status and number offspring, with litter pair as a random effect.

### Modeling the probability of bone failure

Previous research on mechanical properties of rodent femurs found that, among several morphological and compositional traits measured in eight morphologically varying mouse strains, cortical area (CA) at the midshaft was the best predictor of maximum load (defined as the greatest force attained prior to bone failure, measured via four-point bending; published Pearson’s *r* = 0.95) [37]. We therefore used cortical area at the femoral midshaft to predict max load of Damaraland mole-rat femurs. To do so, we first fit a linear model of max load as a function of cortical area (unadjusted for body weight) using published mouse data (R^2^ = 0.877, n = 81, p = 6.64 × 10^−38^) [37]. We extrapolated from this linear fit to predict max load from cortical area at the midshaft of Damaraland mole-rat femurs. Predicted max loads were then used as input for Cox proportional hazards models, comparing either all queens to nonbreeders or queens with ≥ 6 offspring to nonbreeders. Models were fit using the R function *coxph*, and were confirmed to meet the proportional hazards assumption using the *cox.zph* function in the R package *survival* [83]. Because max load was not directly measured in the Damaraland mole-rats, we used the Cox proportional hazards models to specifically evaluate the relative hazard of bone failure depending on queen status/number of offspring. We therefore report the results in Figure 5 based on relative force (with the median predicted failure value for nonbreeders set to 1) instead of absolute force in Newtons.

## Data Availability

All RNA sequencing data generated during this study are available in the NCBI Gene Expression Omnibus (series accession GSE152659). ATAC-Seq data are available in the NCBI Sequence Read Archive (BioProject accession number PRJNA649596). μCT data from this study are available on MorphoSource (http://www.morphosource.org, project 1056).

## Acknowledgements

We thank Tim Vink, Dave Gaynor, and the mole-rat house staff and volunteers for their tremendous contributions to the Kalahari Mole-rat Project. We also thank Irene Garcia, Mari Cobb, Brianna Bowman, Anna Luiza Wolf, Alice Zhou, Yilin Yu, B.J. Nielsen, and Tawni Voyles for their contributions to sample collection and data generation, Graham Treece for guidance on quantifying bone thickness with Stradview, Karl Jepsen for sharing data on mouse femurs, Lou DeFrate for guidance on estimating bone strength, Saideep Gona and Luis Barreiro for their contributions on the footprint analysis, and members of the Tung lab for feedback on earlier versions of this manuscript. Support for this work was provided by the European Research Council (Grants 294494 and 742808 to T.C.B), the Human Frontier Science Program (RGP0051-2017 to J.T., S.M., and T.C.B.), the National Science Foundation (IOS-7801004 to J.T.), the National Institutes of Health (F32HD095616 to R.A.J.), a Sloan Foundation Early Career Research Fellowship to J.T., a Foerster-Bernstein Postdoctoral Fellowship to R.A.J., and a Natural Environmental Research Council Doctoral Training Program to Ja.T. High-performance computing resources were supported by the North Carolina Biotechnology Center (Grant Number 2016-IDG-1013).

## Author Contributions

Conceptualization, R.A.J., T.C.B., and J.T.; Investigation, R.A.J., Ja.T., P.V., H.K., L.S., S.M., C.K., T.C.B., and J.T. Formal Analysis, R.A.J., H.K., S.M.; Writing—Original Draft, R.A.J. and J.T.; Writing—Reviewing & Editing, R.A.J., P.V., Ja.T., H.K., L.S., S.M., C.K., T.C.B., and J.T. Funding Acquisition, J.T. and T.C.B. Supervision, T.C.B. and J.T.

---

**Supplementary Figure S1.**
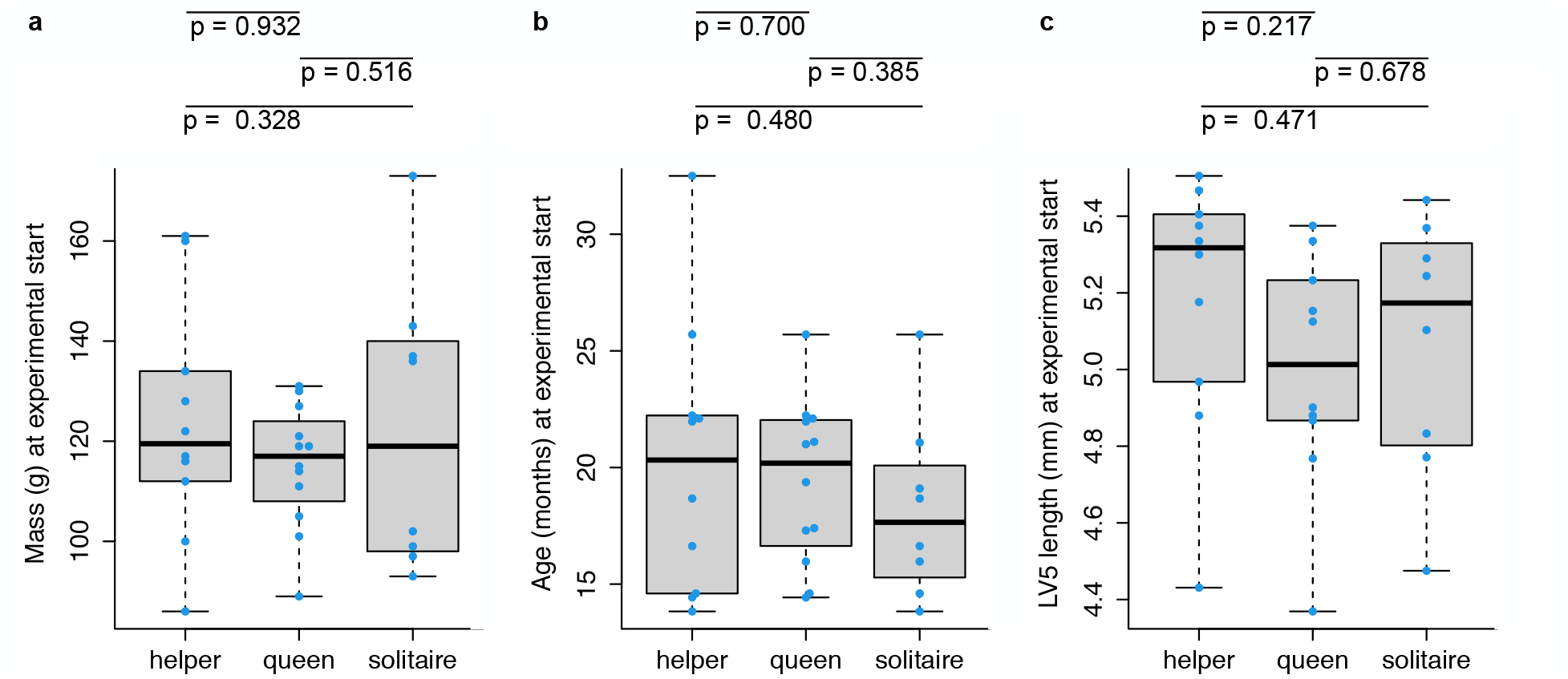
Queen, helper, and solitaire female mole-rats do not differ in mass, age, or lumbar vertebra 5 length at the start of the experiment. Each box represents the interquartile range, with the median value depicted as a horizontal bar. Whiskers extend to the most extreme values within 1.5x of the interquartile range. Dots represent individual animals. P-values are from unpaired t-tests between each pairwise comparison between helpers (n = 10), queens (n = 12), and solitaires (n = 8). For LV5 length, data were only available for 10 queens. Raw data values are provided in Supplementary Table S1.

**Supplementary Figure S2.**
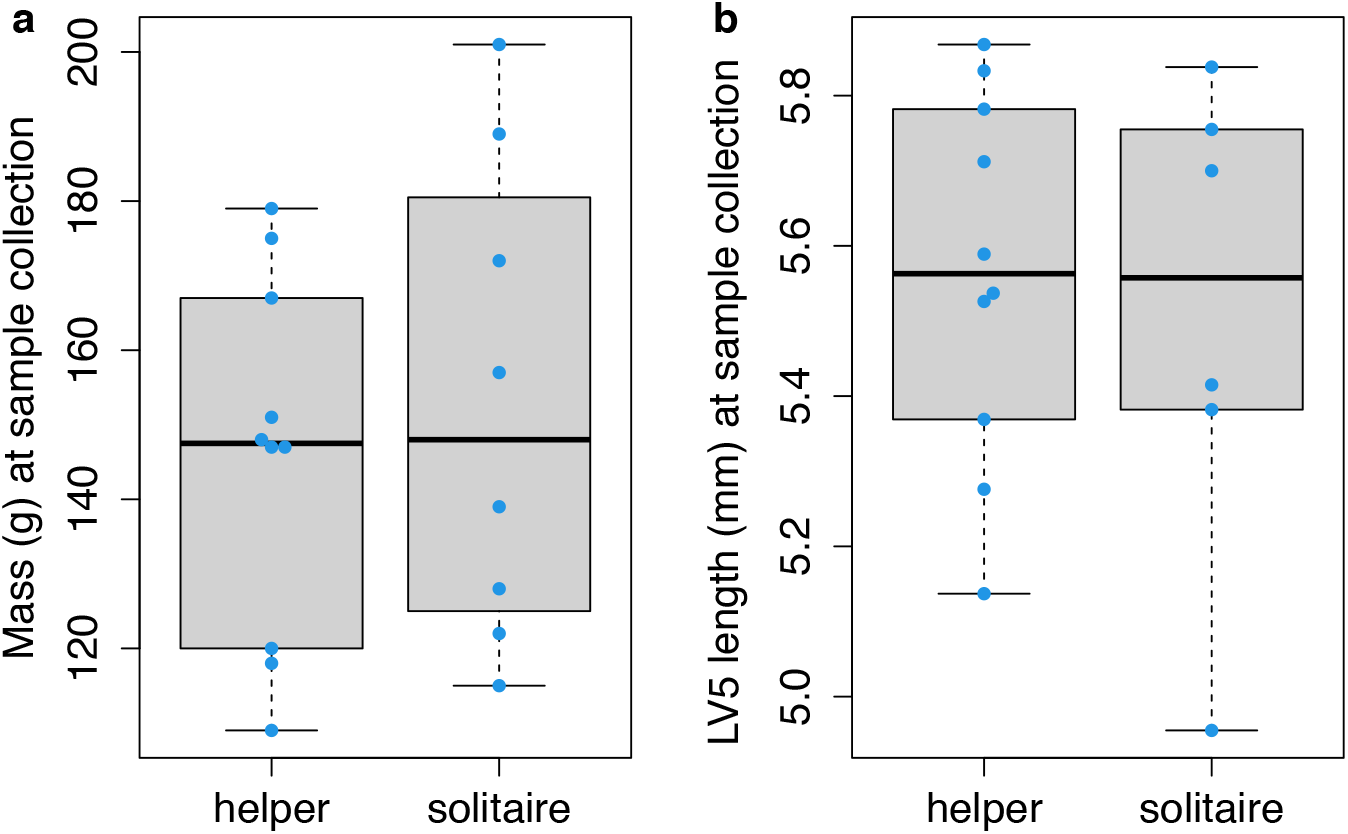
Mass and LV5 length of helper and solitaire female mole-rats after 12 – 22 months of experimental treatment of social status. Helper and solitaire female mole-rats do not differ in (a) mass (unpaired t-test; t = 0.496, df = 12.733, p = 0.629) or (b) lumbar vertebra 5 length (unpaired t-test; t = −0.358, df = 8.391, p = 0.729) after 12 – 22 months of experimental treatment of social status. Each box represents the interquartile range, with the median value depicted as a horizontal bar. Whiskers extend to the most extreme values within 1.5x of the interquartile range. Dots represent individual animals. Raw data values are provided in Supplementary Table S1.

**Supplementary Figure S3.**
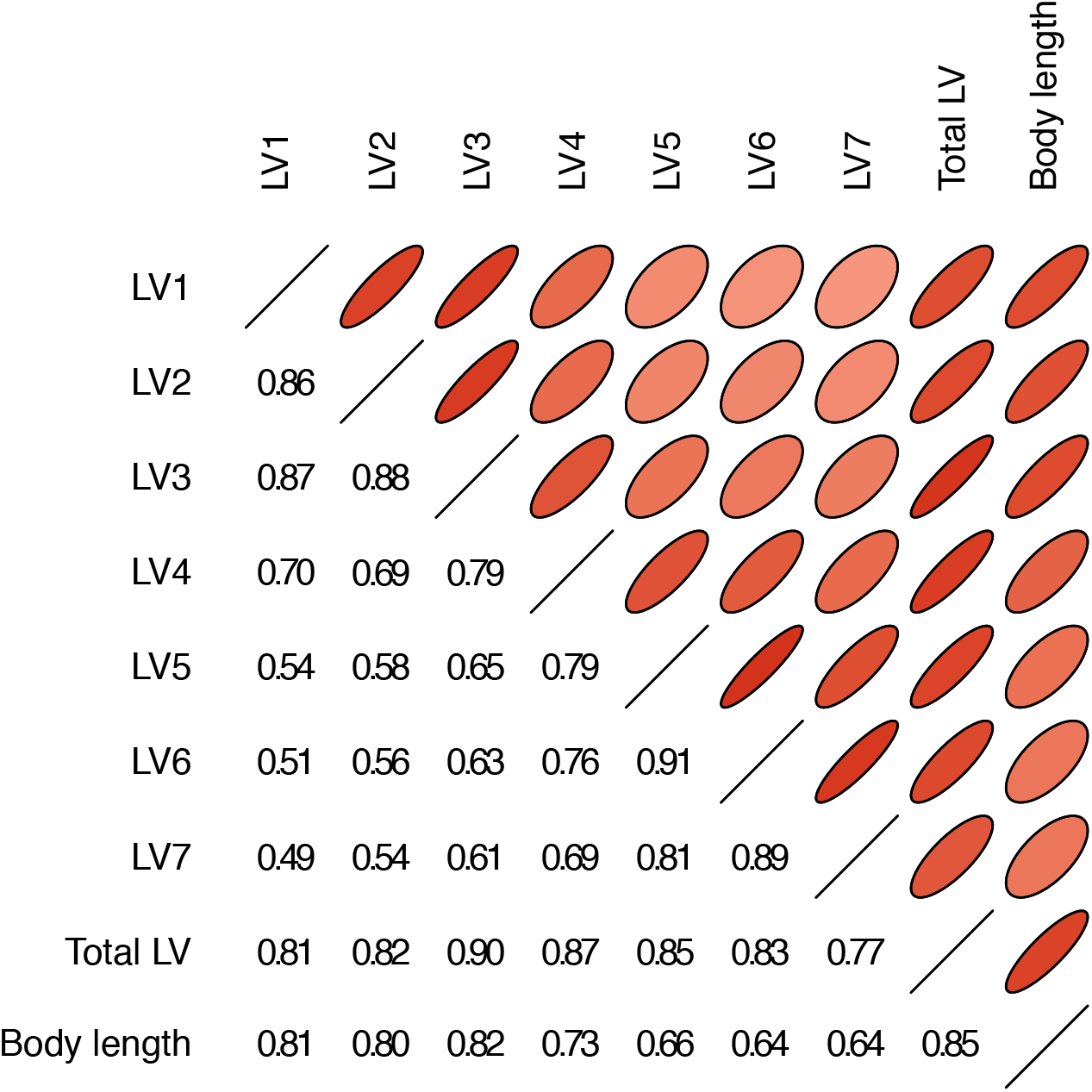
Body length is positively correlated with the lengths of lumbar vertebrae (LV) 1 – 7. Pearson correlations between the length of each lumbar vertebra and body length from mole-rat x-ray data. Narrower ovals with darker shades of red indicate larger Pearson correlations; correlation values are also given in the lower left triangle.

**Supplementary Figure S4.**
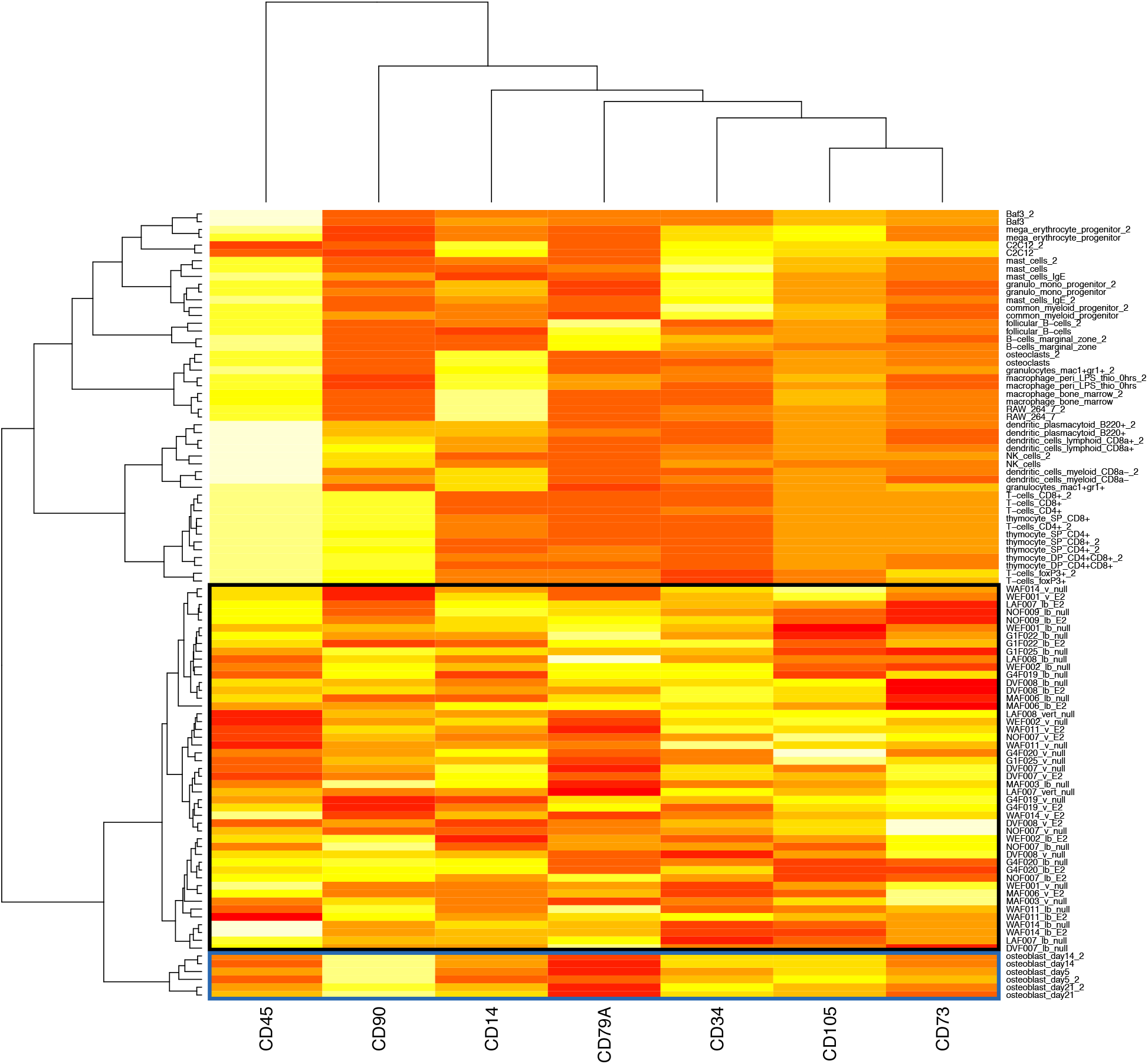
Mole-rat RNA-Seq samples cluster closest with purified mouse osteoblasts, based on canonical bMSC markers. Clustering was performed using Ward’s hierarchical clustering method on Euclidean distances of the quantile normalized expression of the seven bMSC markers (out of 11 described [22]) that were quantified in both the mole-rat and reference mouse [23] data sets. The black box indicates mole-rat samples, and the blue box indicates mouse osteoblasts, a bMSC lineage cell type.

**Supplementary Figure S5.**
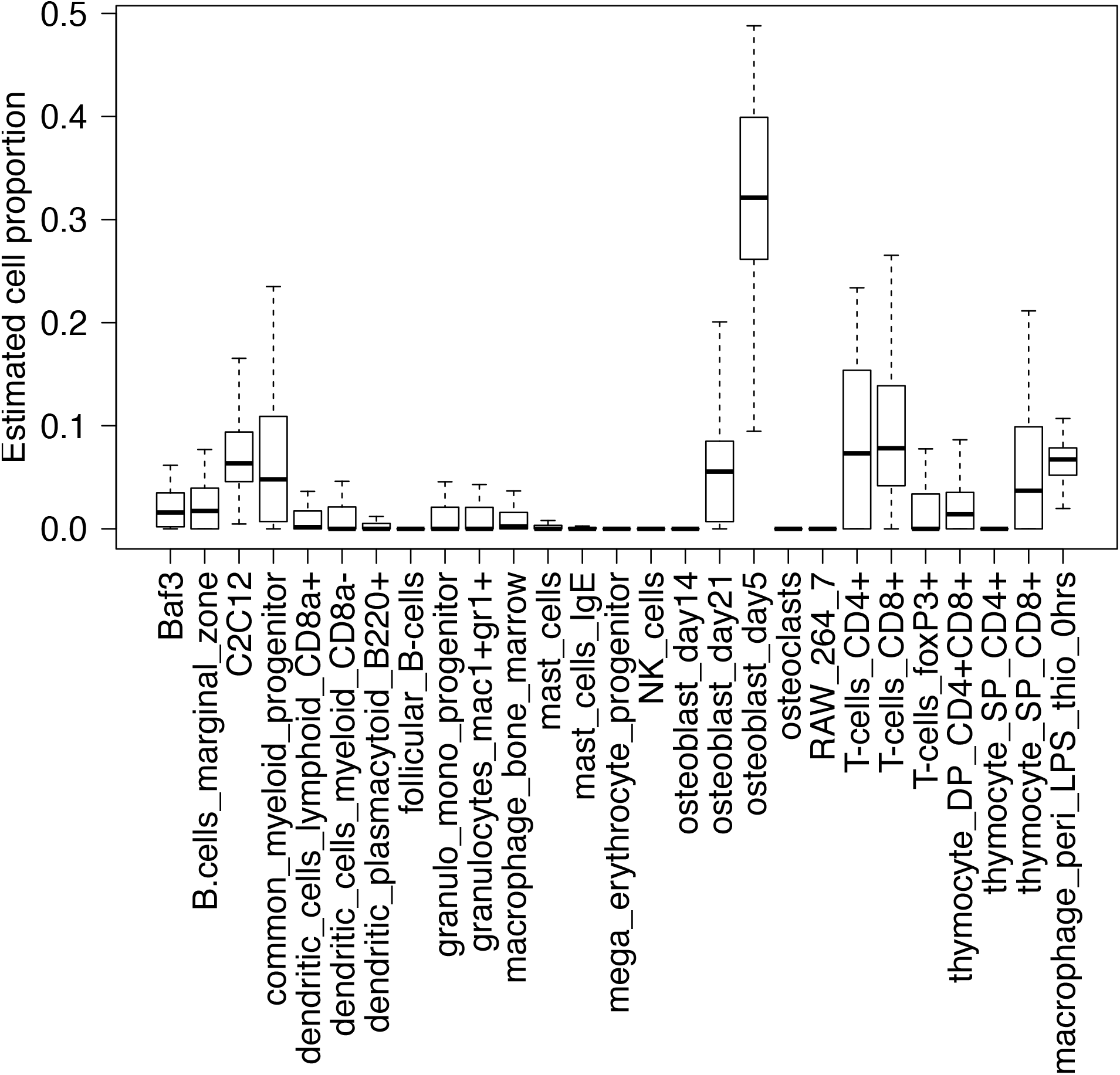
Estimated cell proportions for the 47 mole-rat RNA-Seq samples. Each box represents the interquartile range, with the median value depicted as a horizontal bar. Whiskers extend to the most extreme estimates within 1.5x the interquartile range. Cell proportions were estimated with CIBERSORT [24], based on reference gene expression levels for 412 marker genes in 27 purified mouse cell types [23]. The predicted predominant cell type in the mole-rat samples is most similar to early stage osteoblasts, which are cells from the bMSC lineage.

**Supplementary Figure S6.**
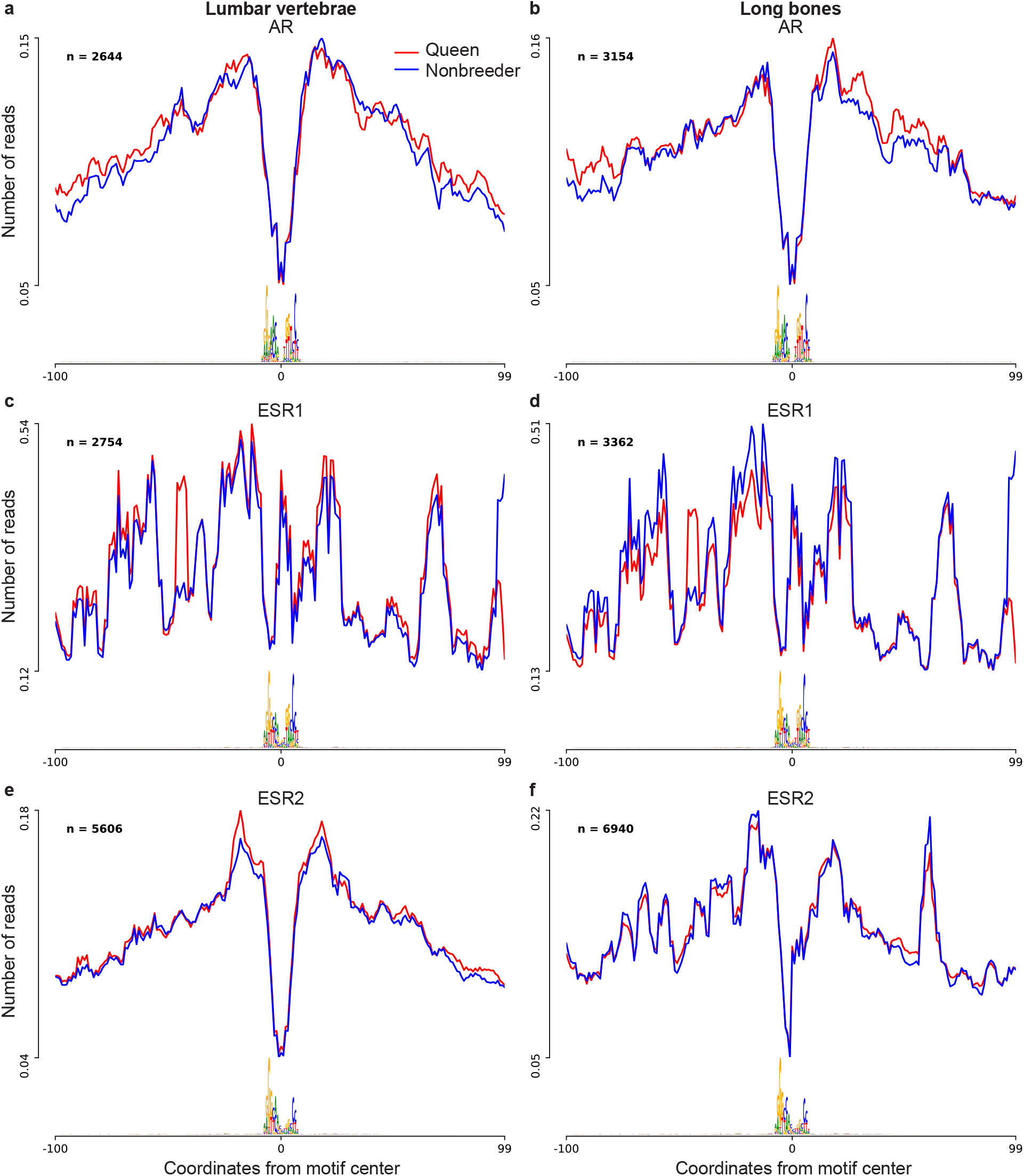
Footprint profiles of transcription factors androgen receptor (AR), estrogen receptor 1 (ESR1), and estrogen receptor 2 (ESR2). Transcription factor footprints were profiled separately for lumbar vertebrae (n = 4; 2 queens and 2 nonbreeders) and for long bones (n = 4; 2 queens and 2 nonbreeders). Transcription factor activity was not significantly different between queens and nonbreeders in any of the three transcription factors, in either bone type (paired t-tests; all p > 0.05).

**Supplementary Figure S7.**
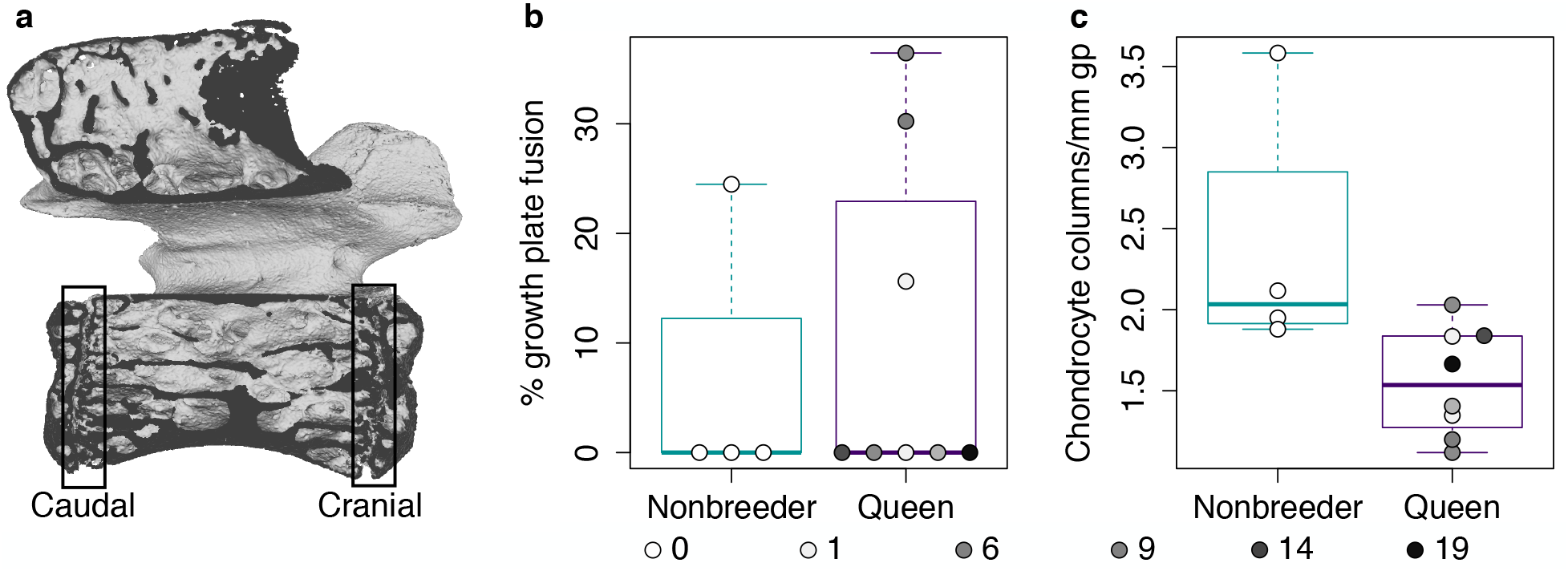
Growth potential in lumbar vertebra 7 (LV7). (a) μCT scan of LV7 of a female Damaraland mole-rat. The boxes indicate the locations of the caudal and cranial growth plates. (b) The number of offspring produced by queens does not significantly predict growth plate fusion (quantified as the average of the caudal and cranial growth plates; β = 2.745 × 10^−4^, p = 0.970, n = 12, controlling for age) or chondrocyte proliferation within the remaining growth plate (β = −0.033, p = 0.293, n = 12, controlling for age). Each box represents the interquartile range, with the median value depicted as a horizontal bar. Whiskers extend to the most extreme values within 1.5x of the interquartile range. Dots represent individual animals, and shading indicates each animal’s total offspring number.

**Supplementary Figure S8.**
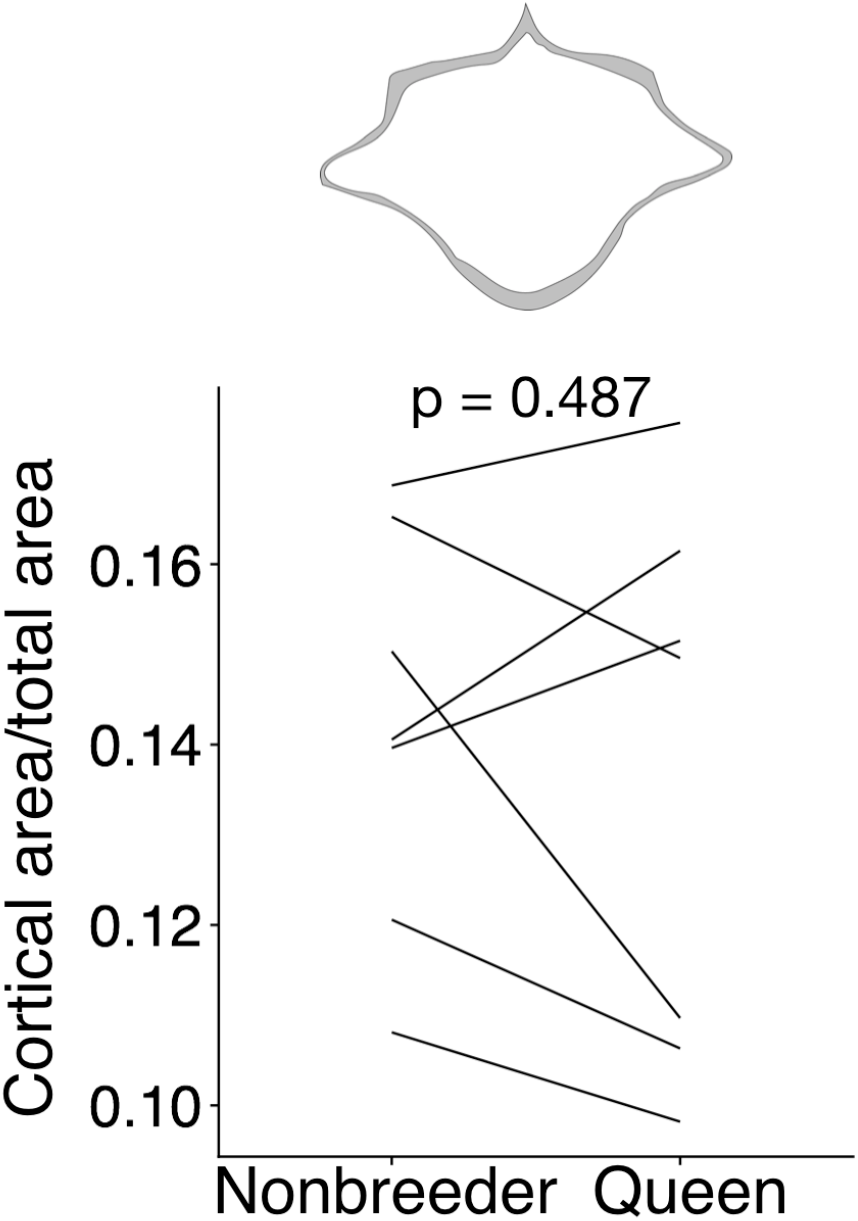
Queens do not exhibit reduced cortical area at the midsection of LV6. At top, cross-section with area highlighted in gray shows the measure represented in the plot. Each line represents an age-matched nonbreeder and queen littermate pair. Queens and nonbreeders show no difference in cortical area/total area at the LV6 midsection (paired t-test of cortical area/total area, t = −0.741, df = 6, p = 0.487).

**Supplementary Figure S9.**
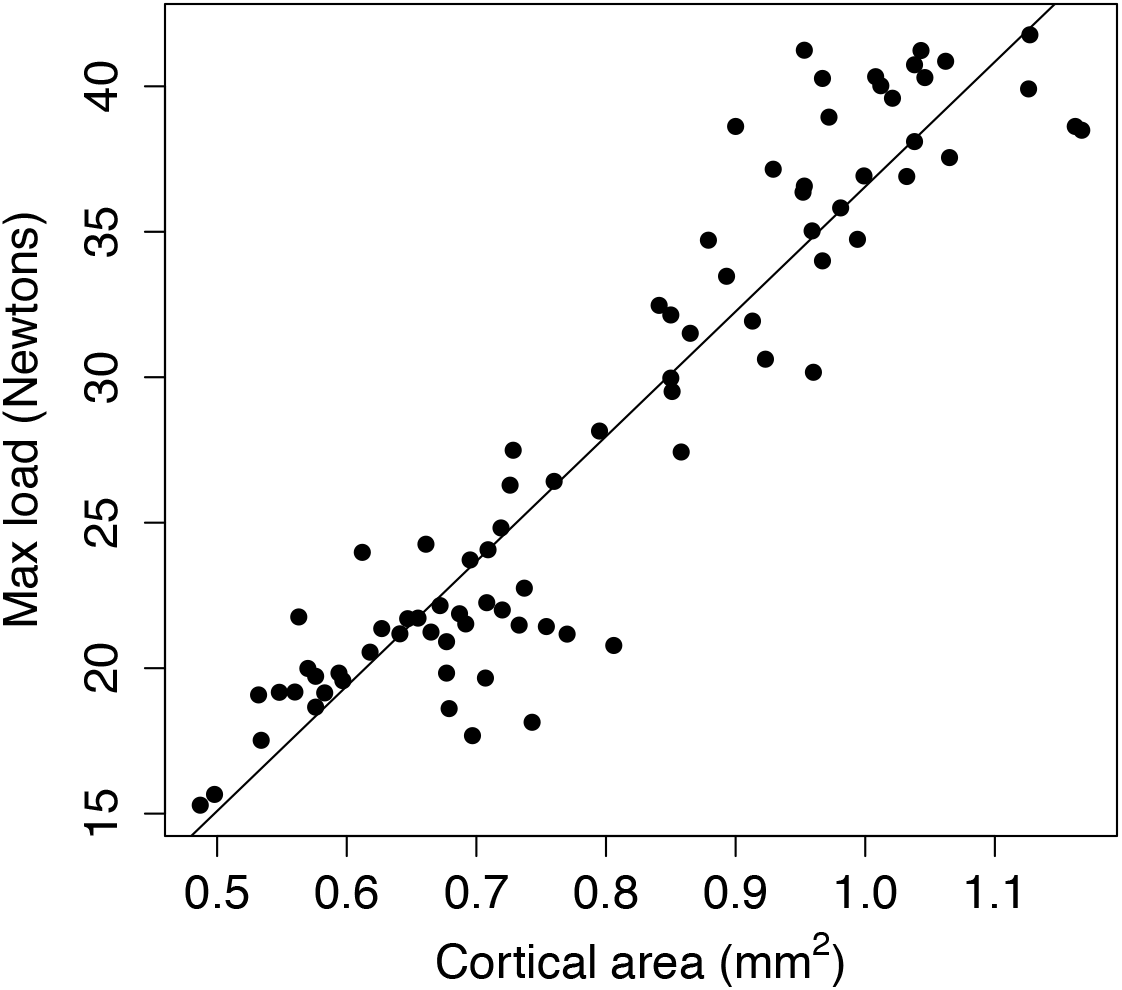
Relationship between max load and cortical area in mouse femurs. Max load shows a highly linear relationship with cortical area across mouse femurs (R^2^ = 0.88, p = 6.64 × 10^−38^). Each dot represents a single mouse femur. Solid line indicates the best fit line. Data are from [37].

## References

1. Wilson, E.O. (1971). The insect societies. The insect societies.

2. Keller, L., and Genoud, M. (1997). Extraordinary lifespans in ants: a test of evolutionary theories of ageing. Nature 389, 958–960.

3. Berndt, K.P., and Eichler, W. (1987). Die Pharaoameise, Monomorium pharaonis (L.)(Hym., Myrmicidae). Mitteilungen aus dem Museum für Naturkunde in Berlin. Zoologisches Museum und Institut für Spezielle Zoologie (Berlin) 63, 3–186.

4. Page Jr, R.E., and Peng, C.Y.-S. (2001). Aging and development in social insects with emphasis on the honey bee, Apis mellifera L. Experimental gerontology 36, 695–711.

5. Smith, C.R., Toth, A.L., Suarez, A.V., and Robinson, G.E. (2008). Genetic and genomic analyses of the division of labour in insect societies. Nature Reviews Genetics 9, 735–748.

6. Koenig, W.D., and Dickinson, J.L. (2016). Cooperative breeding in vertebrates: studies of ecology, evolution, and behavior, (Cambridge University Press).

7. Clutton-Brock, T., Hodge, S., Spong, G., Russell, A., Jordan, N., Bennett, N., Sharpe, L., and Manser, M. (2006). Intrasexual competition and sexual selection in cooperative mammals. Nature 444, 1065–1068.

8. Huchard, E., English, S., Bell, M.B., Thavarajah, N., and Clutton-Brock, T. (2016). Competitive growth in a cooperative mammal. Nature 533, 532–534.

9. O’Riain, M., Jarvis, J., Alexander, R., Buffenstein, R., and Peeters, C. (2000). Morphological castes in a vertebrate. Proceedings of the National Academy of Sciences 97, 13194–13197.

10. Russell, A.F., Carlson, A.A., McIlrath, G.M., Jordan, N.R., and Clutton-Brock, T. (2004). Adaptive size modification by dominant female meerkats. Evolution 58, 1600–1607.

11. Thorley, J., Katlein, N., Goddard, K., Zöttl, M., and Clutton-Brock, T. (2018). Reproduction triggers adaptive increases in body size in female mole-rats. Proceedings of the Royal Society B: Biological Sciences 285, 20180897.

12. Young, A.J., and Bennett, N.C. (2010). Morphological divergence of breeders and helpers in wild Damaraland mole-rat societies. Evolution: International Journal of Organic Evolution 64, 3190–3197.

13. Bens, M., Szafranski, K., Holtze, S., Sahm, A., Groth, M., Kestler, H.A., Hildebrandt, T.B., and Platzer, M. (2018). Naked mole-rat transcriptome signatures of socially suppressed sexual maturation and links of reproduction to aging. BMC biology 16, 77.

14. Mulugeta, E., Marion-Poll, L., Gentien, D., Ganswindt, S.B., Ganswindt, A., Bennett, N.C., Blackburn, E.H., Faulkes, C.G., and Heard, E. (2017). Molecular insights into the pathways underlying naked mole-rat eusociality. bioRxiv, 209932.

15. Sahm, A., Hoffmann, S., Koch, P., Henning, Y., Bens, M., Groth, M., Burda, H., Begall, S., Ting, S., and Goetz, M. (2020). Status-dependent aging rates in long-lived, social mole-rats are shaped by HPA stress axis. bioRxiv.

16. Bennett, N.C., and Faulkes, C.G. (2000). African mole-rats: ecology and eusociality, (Cambridge University Press).

17. Jarvis, J. (1981). Eusociality in a mammal: cooperative breeding in naked mole-rat colonies. Science 212, 571–573.

18. Jarvis, J., and Bennett, N. (1993). Eusociality has evolved independently in two genera of bathyergid mole-rats—but occurs in no other subterranean mammal. Behavioral Ecology and Sociobiology 33, 253–260.

19. Bennett, N., Faulkes, C., and Molteno, A. (1996). Reproductive suppression in subordinate, non-breeding female Damaraland mole-rats: two components to a lifetime of socially induced infertility. Proceedings of the Royal Society of London. Series B: Biological Sciences 263, 1599–1603.

20. Snyman, P., Jackson, C.R., and Bennett, N.C. (2006). Do dispersing non-reproductive female Damaraland mole-rats, Cryptomys damarensis (Rodentia: Bathyergidae) exhibit spontaneous or induced ovulation? Physiology & behavior 87, 88–94.

21. Dengler-Crish, C.M., and Catania, K.C. (2009). Cessation of reproduction-related spine elongation after multiple breeding cycles in female naked mole-rats. The Anatomical Record: Advances in Integrative Anatomy and Evolutionary Biology: Advances in Integrative Anatomy and Evolutionary Biology 292, 131–137.

22. Dominici, M., Le Blanc, K., Mueller, I., Slaper-Cortenbach, I., Marini, F., Krause, D., Deans, R., Keating, A., Prockop, D., and Horwitz, E. (2006). Minimal criteria for defining multipotent mesenchymal stromal cells. The International Society for Cellular Therapy position statement. Cytotherapy 8, 315–317.

23. Hume, D.A., Summers, K.M., Raza, S., Baillie, J.K., and Freeman, T.C. (2010). Functional clustering and lineage markers: insights into cellular differentiation and gene function from large-scale microarray studies of purified primary cell populations. Genomics 95, 328–338.

24. Newman, A.M., Liu, C.L., Green, M.R., Gentles, A.J., Feng, W., Xu, Y., Hoang, C.D., Diehn, M., and Alizadeh, A.A. (2015). Robust enumeration of cell subsets from tissue expression profiles. Nature methods 12, 453.

25. Redlich, K., and Smolen, J.S. (2012). Inflammatory bone loss: pathogenesis and therapeutic intervention. Nature reviews Drug discovery 11, 234–250.

26. Darden, A.G., Ries, W.L., Wolf, W.C., Rodriguiz, R.M., and Key Jr, L.L. (1996). Osteoclastic superoxide production and bone resorption: stimulation and inhibition by modulators of NADPH oxidase. Journal of Bone and Mineral Research 11, 671–675.

27. Datta, H., Rathod, H., Manning, P., Turnbull, Y., and McNeil, C. (1996). Parathyroid hormone induces superoxide anion burst in the osteoclast: evidence for the direct instantaneous activation of the osteoclast by the hormone. Journal of endocrinology 149, 269–275.

28. Key, L., Ries, W., Taylor, R., Hays, B., and Pitzer, B. (1990). Oxygen derived free radicals in osteoclasts: the specificity and location of the nitroblue tetrazolium reaction. Bone 11, 115–119.

29. Key, L., Wolf, W., Gundberg, C., and Ries, W. (1994). Superoxide and bone resorption. Bone 15, 431–436.

30. Segeletz, S., and Hoflack, B. (2016). Proteomic approaches to study osteoclast biology. Proteomics 16, 2545–2556.

31. Bennett, N. (1994). Reproductive suppression in social Cryptomys damarensis colonies—a lifetime of socially-induced sterility in males and females (Rodentia: Bathyergidae). Journal of Zoology 234, 25–39.

32. Crawford, L., Monod, A., Chen, A.X., Mukherjee, S., and Rabadán, R. (2016). Functional data analysis using a topological summary statistic: the smooth Euler characteristic transform. arXiv preprint arXiv:1611.06818.

33. Kilborn, S.H., Trudel, G., and Uhthoff, H. (2002). Review of growth plate closure compared with age at sexual maturity and lifespan in laboratory animals. Journal of the American Association for Laboratory Animal Science 41, 21–26.

34. Kovacs, C.S. (2016). Maternal mineral and bone metabolism during pregnancy, lactation, and post-weaning recovery. Physiological reviews 96, 449–547.

35. Schmidt, C.M., Jarvis, J.U., and Bennett, N.C. (2013). The long-lived queen: reproduction and longevity in female eusocial Damaraland mole-rats (Fukomys damarensis). African Zoology 48, 193–196.

36. Szulc, P., Seeman, E., Duboeuf, F., Sornay-Rendu, E., and Delmas, P.D. (2006). Bone fragility: failure of periosteal apposition to compensate for increased endocortical resorption in postmenopausal women. Journal of bone and mineral research 21, 1856–1863.

37. Jepsen, K.J., Akkus, O.J., Majeska, R.J., and Nadeau, J.H. (2003). Hierarchical relationship between bone traits and mechanical properties in inbred mice. Mammalian Genome 14, 97–104.

38. Henry, E.C., Dengler-Crish, C.M., and Catania, K.C. (2007). Growing out of a caste-reproduction and the making of the queen mole-rat. Journal of Experimental Biology 210, 261–268.

39. Faulkes, C.G., and Bennett, N.C. (2016). Damaraland and naked mole-rats: convergence of social evolution. Cooperative breeding in vertebrates, 338–352.

40. Cutler Jr, G. (1997). The role of estrogen in bone growth and maturation during childhood and adolescence. The Journal of steroid biochemistry and molecular biology 61, 141–144.

41. Khalid, A.B., and Krum, S.A. (2016). Estrogen receptors alpha and beta in bone. Bone 87, 130–135.

42. Juul, A. (2001). The effects of oestrogens on linear bone growth. Apmis 109, S124–S134.

43. Houslay, T.M., Vullioud, P., Zöttl, M., and Clutton-Brock, T.H. (2020). Benefits of cooperation in captive Damaraland mole-rats. Behavioral Ecology.

44. Clutton-Brock, T.H., and Manser, M. (2016). Meerkats: cooperative breeding in the Kalahari. Cooperative breeding in vertebrates 294, 317.

45. Morseth, B., Emaus, N., and Jørgensen, L. (2011). Physical activity and bone: The importance of the various mechanical stimuli for bone mineral density. A review. Norsk epidemiologi 20.

46. Jarvis, J.U., O’Riain, M.J., Bennett, N.C., and Sherman, P.W. (1994). Mammalian eusociality: a family affair. Trends in Ecology & Evolution 9, 47–51.

47. Pinto, M., Jepsen, K., Terranova, C., and Buffenstein, R. (2010). Lack of sexual dimorphism in femora of the eusocial and hypogonadic naked mole-rat: a novel animal model for the study of delayed puberty on the skeletal system. Bone 46, 112–120.

48. Burda, H., Honeycutt, R.L., Begall, S., Locker-Grütjen, O., and Scharff, A. (2000). Are naked and common mole-rats eusocial and if so, why? Behavioral Ecology and Sociobiology 47, 293–303.

49. Rodrigues, M.A., and Flatt, T. (2016). Endocrine uncoupling of the trade-off between reproduction and somatic maintenance in eusocial insects. Current opinion in insect science 16, 1–8.

50. Rueppell, O., Aumer, D., and Moritz, R.F. (2016). Ties between ageing plasticity and reproductive physiology in honey bees (Apis mellifera) reveal a positive relation between fecundity and longevity as consequence of advanced social evolution. Current opinion in insect science 16, 64–68.

51. Zöttl, M., Vullioud, P., Mendonça, R., Ticó, M.T., Gaynor, D., Mitchell, A., and Clutton-Brock, T. (2016). Differences in cooperative behavior among Damaraland mole rats are consequences of an age-related polyethism. Proceedings of the National Academy of Sciences 113, 10382–10387.

52. Schneider, C.A., Rasband, W.S., and Eliceiri, K.W. (2012). NIH Image to ImageJ: 25 years of image analysis. Nature methods 9, 671–675.

53. Martin, M. (2011). Cutadapt removes adapter sequences from high-throughput sequencing reads. EMBnet. journal 17, 10–12.

54. Fang, X., Seim, I., Huang, Z., Gerashchenko, M.V., Xiong, Z., Turanov, A.A., Zhu, Y., Lobanov, A.V., Fan, D., and Yim, S.H. (2014). Adaptations to a subterranean environment and longevity revealed by the analysis of mole rat genomes. Cell reports 8, 1354–1364.

55. Dobin, A., Davis, C.A., Schlesinger, F., Drenkow, J., Zaleski, C., Jha, S., Batut, P., Chaisson, M., and Gingeras, T.R. (2013). STAR: ultrafast universal RNA-seq aligner. Bioinformatics 29, 15–21.

56. Anders, S., Pyl, P.T., and Huber, W. (2014). HTSeq–A Python framework to work with high-throughput sequencing data. Bioinformatics, btu638.

57. Wagner, G.P., Kin, K., and Lynch, V.J. (2012). Measurement of mRNA abundance using RNA-seq data: RPKM measure is inconsistent among samples. Theory in biosciences 131, 281–285.

58. Law, C.W., Chen, Y., Shi, W., and Smyth, G.K. (2014). Voom: precision weights unlock linear model analysis tools for RNA-seq read counts. Genome biology 15, R29.

59. Robinson, M.D., and Oshlack, A. (2010). A scaling normalization method for differential expression analysis of RNA-seq data. Genome biology 11, R25.

60. Anders, S., and Huber, W. (2010). Differential expression analysis for sequence count data. Genome Biol 11, R106.

61. Smyth, G.K. (2005). Limma: linear models for microarray data. In Bioinformatics and computational biology solutions using R and Bioconductor. (Springer), pp. 397–420.

62. Akdemir, D., and Okeke, U. (2015). EMMREML: Fitting mixed models with known covariance structures. R package version 3.

63. Storey, J.D., and Tibshirani, R. (2003). Statistical significance for genomewide studies. Proc Natl Acad Sci U S A 100, 9440–9445.

64. Raudvere, U., Kolberg, L., Kuzmin, I., Arak, T., Adler, P., Peterson, H., and Vilo, J. (2019). g: Profiler: a web server for functional enrichment analysis and conversions of gene lists (2019 update). Nucleic acids research 47, W191–W198.

65. McKenna, A., Hanna, M., Banks, E., Sivachenko, A., Cibulskis, K., Kernytsky, A., Garimella, K., Altshuler, D., Gabriel, S., and Daly, M. (2010). The Genome Analysis Toolkit: a MapReduce framework for analyzing next-generation DNA sequencing data. Genome research 20, 1297–1303.

66. Danecek, P., Auton, A., Abecasis, G., Albers, C.A., Banks, E., DePristo, M.A., Handsaker, R.E., Lunter, G., Marth, G.T., and Sherry, S.T. (2011). The variant call format and VCFtools. Bioinformatics 27, 2156–2158.

67. Browning, S.R., and Browning, B.L. (2007). Rapid and accurate haplotype phasing and missing-data inference for whole-genome association studies by use of localized haplotype clustering. The American Journal of Human Genetics 81, 1084–1097.

68. Phinney, D.G., Kopen, G., Isaacson, R.L., and Prockop, D.J. (1999). Plastic adherent stromal cells from the bone marrow of commonly used strains of inbred mice: variations in yield, growth, and differentiation. Journal of cellular biochemistry 72, 570–585.

69. Buenrostro, J.D., Giresi, P.G., Zaba, L.C., Chang, H.Y., and Greenleaf, W.J. (2013). Transposition of native chromatin for fast and sensitive epigenomic profiling of open chromatin, DNA-binding proteins and nucleosome position. Nature methods 10, 1213.

70. Corces, M.R., Trevino, A.E., Hamilton, E.G., Greenside, P.G., Sinnott-Armstrong, N.A., Vesuna, S., Satpathy, A.T., Rubin, A.J., Montine, K.S., and Wu, B. (2017). An improved ATAC-seq protocol reduces background and enables interrogation of frozen tissues. Nature methods 14, 959.

71. Krueger, F. (2015). Trim Galore. http://www.bioinformatics.babraham.ac.uk/projects/trim_galore/.

72. Li, H., and Durbin, R. (2010). Fast and accurate long-read alignment with Burrows– Wheeler transform. Bioinformatics 26, 589–595.

73. Zhang, Y., Liu, T., Meyer, C.A., Eeckhoute, J., Johnson, D.S., Bernstein, B.E., Nusbaum, C., Myers, R.M., Brown, M., and Li, W. (2008). Model-based analysis of ChIP-Seq (MACS). Genome biology 9, R137.

74. Heinz, S., Benner, C., Spann, N., Bertolino, E., Lin, Y.C., Laslo, P., Cheng, J.X., Murre, C., Singh, H., and Glass, C.K. (2010). Simple combinations of lineage-determining transcription factors prime cis-regulatory elements required for macrophage and B cell identities. Molecular cell 38, 576–589.

75. Li, Z., Schulz, M.H., Look, T., Begemann, M., Zenke, M., and Costa, I.G. (2019). Identification of transcription factor binding sites using ATAC-seq. Genome biology 20, 45.

76. Quinlan, A.R., and Hall, I.M. (2010). BEDTools: a flexible suite of utilities for comparing genomic features. Bioinformatics 26, 841–842.

77. Sandelin, A., Alkema, W., Engström, P., Wasserman, W.W., and Lenhard, B. (2004). JASPAR: an open-access database for eukaryotic transcription factor binding profiles. Nucleic acids research 32, D91–D94.

78. Boyer, D.M., Puente, J., Gladman, J.T., Glynn, C., Mukherjee, S., Yapuncich, G.S., and Daubechies, I. (2015). A new fully automated approach for aligning and comparing shapes. The Anatomical Record 298, 249–276.

79. Dimitriadou, E., Hornik, K., Leisch, F., Meyer, D., and Weingessel, A. (2008). Misc functions of the Department of Statistics (e1071), TU Wien. R package 1, 5–24.

80. Golland, P., Liang, F., Mukherjee, S., and Panchenko, D. (2005). Permutation tests for classification. International Conference on Computational Learning Theory, 501–515.

81. Treece, G.M. (2019). http://mi.eng.cam.ac.uk/~gmt11/stradview.

82. Treece, G.M., Gee, A.H., Mayhew, P., and Poole, K.E. (2010). High resolution cortical bone thickness measurement from clinical CT data. Medical image analysis 14, 276–290.

83. Therneau, T.M. (2020). A Package for Survival Analysis in R. R package version 3.1–12, available at https://CRAN.R-project.org/package=survival.

